# Single-cell approach dissecting *agr* quorum sensing dynamics in *Staphylococcus aureus*

**DOI:** 10.1101/2024.02.27.582246

**Authors:** Julian Bär, Samuel G. V. Charlton, Andrea Tarnutzer, Giovanni Stefano Ugolini, Eleonora Secchi, Annelies S. Zinkernagel

**Affiliations:** Department of Infectious Diseases and Hospital Epidemiology, University Hospital Zürich, University of Zürich, Zürich, Switzerland; Institute of Environmental Engineering, ETH Zürich, Zürich, Switzerland

**Keywords:** *Staphylococcus aureus*, *agr*, Quorum sensing, AIP sensitivity, Intergenerational stability, Community interaction, Microfluidics, Fluorescence microscopy, Deep learning, Image analysis

## Abstract

*Staphylococcus aureus* both colonizes humans and causes severe virulent infections. Virulence is regulated by the *agr* quorum sensing system and its autoinducing peptide (AIP), with dynamics at the single-cell level across four *agr*-types – each defined by distinct AIP sequences and capable of cross-inhibition – remaining elusive. Employing microfluidics, time-lapse microscopy, and deep-learning image analysis, we uncovered significant differences in AIP sensitivity among *agr*-types. We observed bimodal *agr* activation, attributed to intergenerational phenotypic stability and influenced by AIP concentration. Upon AIP stimulation, *agr-III* showed AIP insensitivity, while *agr-II* exhibited increased sensitivity and prolonged generation time. Beyond expected cross-inhibition of *agr-I* by heterologous AIP-II and -III, the presumably cross-activating AIP-IV also inhibited *agr-I*. Community interactions across different *agr*-type pairings revealed four main patterns: stable or switched dominance, and delayed or stable dual activation, influenced by community characteristics. These insights underscore the potential of personalized treatment strategies considering virulence and genetic diversity.

## Introduction

*Staphylococcus aureus* colonizes up to 50% of healthy individuals^1^. However, it can also cause a wide array of infections ranging from soft tissue to life-threatening cardiovascular and prosthetic joint infections^2^. Its virulence factor production, often inactive during colonization and chronic infections characterized by biofilm formation^3,4^ or antibiotic persistence^5^, becomes activated in acute, invasive diseases. The *accessory gene regulator* (*agr*) system plays a pivotal role in this virulence regulation^6,7^. Mutations in the *agr* system were linked to less severe disease outcomes^6,8^, such as reduced mortality from pneumonia and decreased necrosis in skin infections in a mouse model^9^. Thus, dissecting the mechanisms behind the ability of *S. aureus* to regulate its virulence expression is vital for preventing its shift from a harmless commensal to a life-threatening pathogen.

The *agr* system is a quorum sensing (QS) system and is regulated via secreted autoinducing peptides (AIPs). Increased extracellular AIP concentrations correlate with high population density and/or restricted diffusion^7,10^. The *agr* system consists of two transcripts, RNAII and RNAIII, which are controlled by the promoters P2 and P3^6^. The RNAII operon encompasses all the genes required for AIP production, secretion, and sensing while RNAIII is the downstream small RNA effector regulating virulence gene expression. AgrA, the response regulator encoded on RNAII, activates P2, driving a positive feedback loop of upregulated RNAII expression leading to increased AIP secretion. At high extracellular AIP concentrations, AgrA activates additionally P3. Activation of the *agr* system, indicated as P3-RNAIII production, upregulates virulence gene expression and is associated with a switch from an avirulent biofilm-related to a virulent dispersal-related gene expression profile^6,11^.

Four *S. aureus agr-*types are known, and each is characterized by a unique AIP amino acid sequence (7 to 9 aa) and correlated with specific combinations of toxin-mediated diseases^12^. The *agr-*types show strong AIP specificity as heterologous AIP inhibits *agr* activity^13,14^. However, moderate cross-activation between *agr*-type *I* and *IV* is reported, likely due to just a single amino acid difference of the respective AIP sequences^13^.

Understanding QS at the single-cell level offers critical insights into bacterial communication that bulk experiments cannot capture. QS heterogeneity or bimodality has been documented, indicating a mix of QS-activated and -inactivated states within bacterial populations^11,15,16^. However, these observations were typically derived indirectly using bulk cultures – either grown on agar or in liquid media – where QS signals could accumulate or establish spatial gradients, which can mask the dynamics of QS by allowing bacteria to experience different local conditions. The physiological state of the bacteria and the environmental factors like pH and nutrient levels further influence QS^16,17^, complicating the interpretation of heterogeneity as an intrinsic QS feature or a byproduct of external conditions.

The ability of *S. aureus* to invade established communities – and potentially cause infections – may be linked to the sensitivity of the *agr* system to interfering stimuli. Notably, commensal species, including other staphylococci producing a variety of AIPs, are known to interfere with *S. aureus* growth and *agr* expression^18–21^. Despite this, tools for studying community interactions at the single-cell level have been scarce^22,23^, leaving a gap in our understanding of how different *agr*-types respond when challenged by a competing strain or species producing heterologous AIPs. Our study addresses this gap, providing unique insights into the spatially structured dynamics of *agr* interactions in a competitive environment.

We unraveled the *agr* response-time and AIP sensitivity profiles of the four known *agr-* types in *S. aureus* at the single-cell level. Mother-machine microfluidics and synthetic AIPs enabled us to study QS heterogeneity in a chemically stable and spatially homogenous environment and further investigate the link between QS and bacterial physiology. Next, we combined heterologous and homologous AIPs, to study *agr* cross-inhibition attributable exclusively to the AIPs at the single-cell level allowing nuanced interpretation of published bulk experiments. Finally, we focused on community interaction dynamics among the four *agr-*types using a novel microfluidic chip design to study crosstalk effects of secreted molecules, including AIPs.

## Results

### Differential AIP sensitivity between *agr-*types

To investigate differences in AIP sensitivity among the four *agr-*types, we utilized microfluidics to switch between growth media with and without synthetic, homologous AIP (Fig.1A) coupled to mother-machine devices containing main medium flow channels and perpendicular side-chambers in which bacteria are embedded in a single-cell layer (Fig.1B). QS activity was monitored using fluorescence microscopy of a P3-YFP reporter, i.e., bacteria were only fluorescent when their *agr* system was activated (Fig.1C, Suppl. Movies1). We validated the reliability of our reporter strains and AIPs using a plate reader (Suppl. Fig.1).

**Figure 1:**
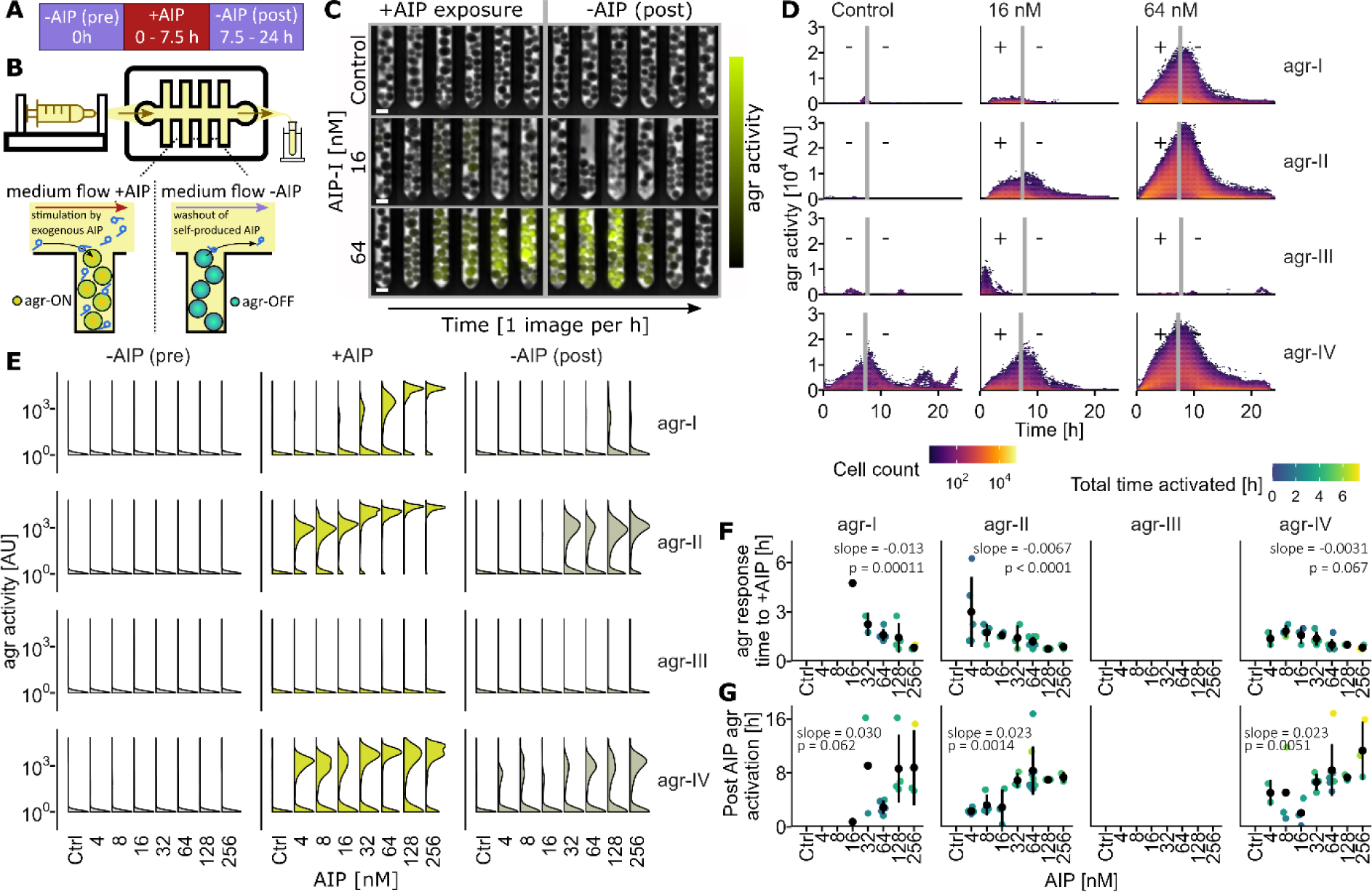
AIP sensitivity differences between *S. aureus agr-*types. **A.** Bacteria were transiently exposed for 7.5 h to synthetic, homologous AIP in mother-machine microfluidic chips. Each experiment was repeated at least three times. **B.** Schematic of mother-machine chip (top) and embedded bacteria (bottom). Growth medium (TSB) was provided at 0.5 ml/h using a syringe pump. This ensured self-produced AIP washout, allowing only exogenous *agr* stimulation. *agr* activity was measured using fluorescence of a P3-YFP reporter plasmid. **C.** Representative image sequences of *Staphylococcus aureus agr-I* in mother-machine devices exposed to three AIP-I concentrations. Phase-contrast images are overlaid with the YFP channel. The vertical gray line indicates AIP-I removal. Contrast adjustments have been performed for visualization purposes. Scale bar = 5 µm. **D.** Single-cell *agr* activity during AIP exposure (concentration annotated above each column) and after AIP removal (indicated by gray lines and plus/minus symbols). Each row represents one *agr-*type. Dot color reflects log_10_ cell counts. On average, 1448728 cells per condition were imaged (details in Suppl. Table1). **E.** Histograms depict *agr* activity of bacteria observed during specific timespans. -AIP (pre): 2 to 0 h before exposure start, +AIP: the final 2 h of exposure, -AIP (post): 3 to 5 h post AIP removal. **F.** Response time to AIP stimulation, quantified as the time when more than 50% of bacteria were activated. **G.** Duration of *agr* activation after removal of AIP stimulation, quantified as the time when less than 50% of bacteria were activated. **FG.** Each dot represents a biological replicate and is colored by total *agr* activation time. Black dots and bars show mean and standard deviation. Statistical significance was assessed with linear regression using log_2_ AIP concentration (slope parameter given for this transformation) followed by estimated marginal means *post-hoc* tests. Only conditions/replicates in which the threshold of 50% activated cells was reached are shown.

We observed strong AIP sensitivity differences among the four *agr*-types (Fig.1DE). No *agr* activity was observed without external AIP stimulation for *agr-*types *I*, *II*, and *III* but occasional low-level *agr* activity was detected for *agr-IV* (Fig.1D, Control). This low-level *agr-IV* activity stemmed from a small subpopulation (maximal proportion of activated bacteria, average: 5.51%, invisible on the scale of Fig.1E, see Suppl. Fig.2). While *agr-III* remained unresponsive to homologous AIP, all other *agr-*types displayed a dose-dependent increase in *agr* activity upon AIP stimulation (Fig.1D).

**Figure 2:**
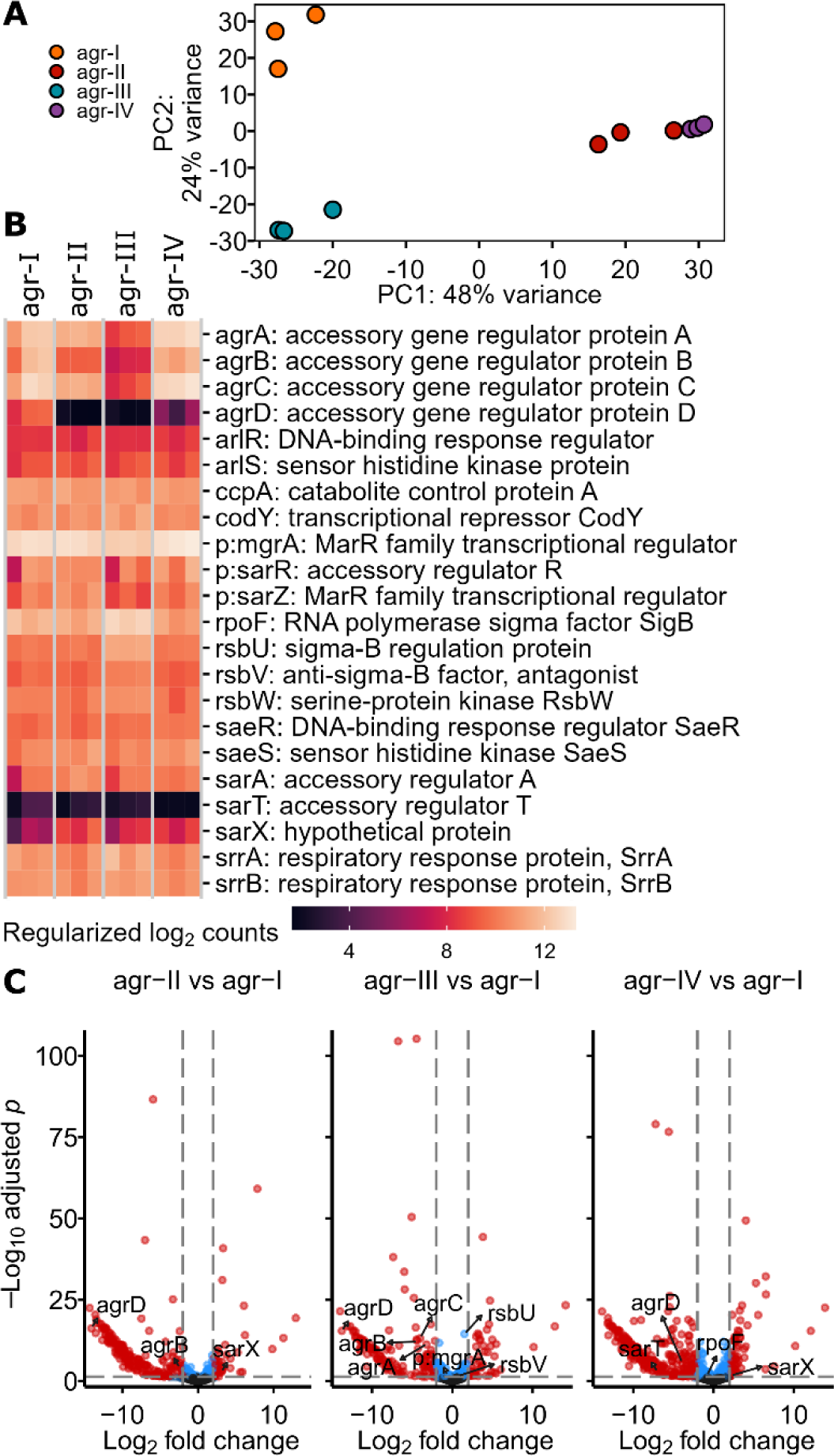
Minimal differences in baseline expression of regulators interacting with the *agr* system. RNA sequencing of exponentially growing cultures without AIP stimulation was performed to quantify gene expression profiles of the four *agr-*types to investigate if these could explain AIP sensitivity differences observed in the mother-machine experiments. **A.** Principal component (PC) analysis of regularized log_2_ (rlog) transformed RNA sequencing counts. Dots represent three independent biological replicates. PC1 and PC2 explained 48% and 24% of the variance, respectively. Axis ratio is scaled accordingly. **B.** Heatmap of rlog transformed RNA counts of the four genes of the RNAII operon and regulators known to interact with the *agr* system. Columns represent *agr-*type and biological replicates. Rows were sorted according to hierarchical clustering. Gene names starting with “p:” originate from the pan genome annotation of AureoWiki. **C.** Volcano plots of DESeq2 based pairwise comparisons of any *agr-*type with *agr-I*. Genes shown in B which had either an adjusted p-value smaller than 0.05 and/or an absolute log_2_ fold change bigger than 2 are annotated.

The distribution of *agr* activity was bimodal, indicating QS heterogeneity (Fig.1E). The ratio of peak heights in the bimodal distribution was dependent on both *agr*-type and AIP concentration, with a larger fraction of bacteria in the second peak at higher AIP levels. *agr-II* was close to full activation at concentrations of 16 nM or higher (maximal proportion of activated bacteria, average: 96.8%, Suppl. Fig.2B), while *agr-I* and *agr-IV* retained a subpopulation in an *agr* inactivated state across all tested concentrations (Fig.1E, Suppl. Fig.2B, Suppl. Animation1).

Clear differences in sensitivity and response time to AIP stimulation (or removal thereof) among *agr-*types were evident. Both *agr-II* and *IV* displayed rapid *agr* activation and at 4-fold lower concentrations than *agr-I* (Fig.1F). Raising AIP concentration above the *agr* activation thresholds significantly shortened the activation time for *agr-I* and *II*, but not for *agr-IV* (Fig.1F) – which likely represented maximal adaptation speed of this *agr*-type – and led to a significantly prolonged activated state for *agr-II* and *IV* (Fig.1G). Though *agr-I* exhibited a similar trend, it lacked statistical significance, probably due to lower number of replicates reaching the threshold of 50% activation. Further, subpopulations of *agr-II* and *IV* retained their activated state until the experiment ended (Suppl. Fig.2B, Suppl. Animation1).

### AIP sensitivity differences are not explained by differential expression of *agr* regulators

We hypothesized that the differences in AIP sensitivity among the *agr*-types might stem from differential gene expression profiles of regulators influencing the *agr* system. To explore this, we conducted RNA sequencing of exponentially growing cultures mimicking the conditions in the microfluidic experiments. Principal component analysis revealed that *agr-II* and *agr-IV* formed a cluster, clearly separated from the other two *agr*-types (Fig.2A). The *agr-III* strain had significantly lower baseline expression of all four genes of the RNAII operon (*agrBDCA,* log2 fold change (LFC) < -2, Fig.2BC). The AIP precursor is encoded on *agrD*, and the differences in the sequences of this gene in *agr-II*, *-III* and *-IV* may negatively impact alignment to the reference genome (USA300_FPR3757, *agr-I*). Therefore, the detected differences in *agrD* expression were likely confounded by reduced alignment quality.

Gene expression of known *agr* regulators was very similar between the four *agr*-types (Fig.2BC). There were two significant differentially expressed genes with absolute LFC larger than 2: Compared to *agr-I*, *sarX* (P2 and P3 repressor^24^) expression was increased in *agr-*types *II* and *IV* (LFC 3.08 and 2.19, respectively) and *sarT* (P3 repressor^25^) was downregulated in *agr-IV* (LFC-6.95). Adjusted p-values of *mgrA* (positive *agr* regulator^6,26^), *rsbU* and *rsbV* (positive regulators of the alternative sigma factor B^27^ which represses P3 expression^6,28^) indicate differential expression of these in *agr-III*, however the absolute LFC were below the threshold of 2 (Fig.2C).

The four *agr*-types had a core genome comprising 2132 genes (62.4% of genes) and 3.6% to 9.7% unique genes (Suppl. Fig.3). DESeq2 identified 221, 175 and 305 significantly differentially expressed genes of *agr-II, agr-III* and *agr-IV* compared to *agr-I*, respectively, which confirmed that genetic background had a strong influence on gene expression profile (Suppl. Fig.4, Suppl. Data1).

**Figure 3:**
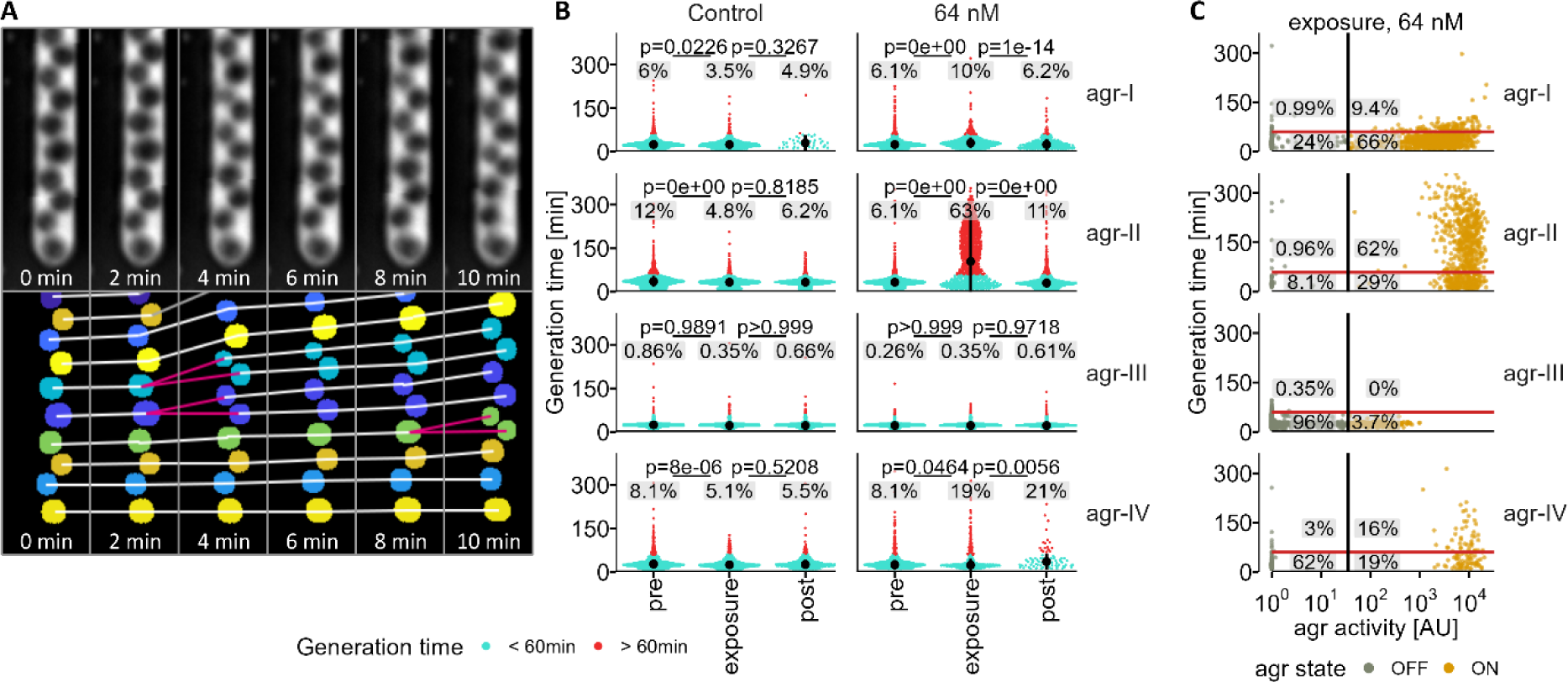
Increased generation time is linked with an activated *agr* system. **A.** Representative image sequence of single-cell segmentation and tracking. Same cells across frames are marked with same color and connected with white lines. Pink lines mark division events. Each experiment was performed three to five times. **B.** Generation time, measured as time between division events, per *agr-*type and with or without AIP exposure. Generation time was measured during predefined experimental phases. pre: 2 to 0 h before AIP exposure; exposure: last 2 h of AIP exposure; post: 3 to 5 h after AIP removal. Colored dots represent individual bacteria with a generation time shorter or longer than 60 min indicated by blue or red, respectively. Percentages of bacteria with prolonged generation times are annotated. Black dots and bars represent median and interquartile range. Statistical significance was assessed with a mixed effect model on log_10_ transformed generation time followed by estimated marginal means *post-hoc* tests. Number of cells analyzed for the four *agr-*types: 7446, 12071, 29645 and 5327. **C.** Bacteria were classified based on generation time (threshold 60 min, red horizontal line) and *agr* activity (threshold 35 AU, black vertical line) with percentages calculated for each quadrant defined by the dual thresholds. Dot color represents *agr* state.

**Figure 4:**
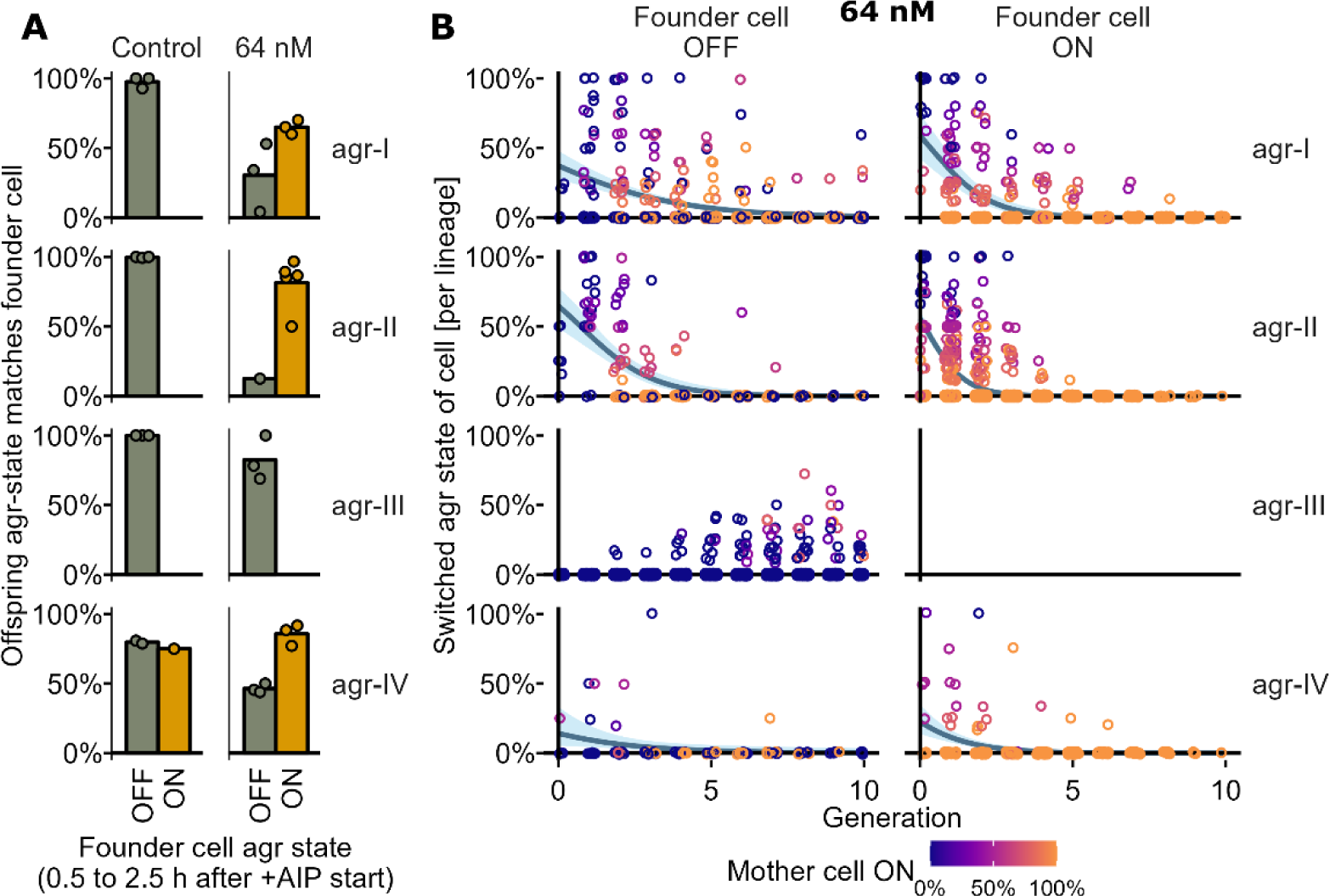
Intergenerational stability of *agr* state. **A.** Through cell tracking, we mapped lineage trees, which represent all offspring from a single founder cell. The x-axis categorizes lineages by the founder cell *agr* state. Founder cells were defined as bacteria which divided during the timeframe of 0.5 to 2.5 h after AIP exposure started. All offspring of these cells were again categorized according to the *agr* state (included up until the end of AIP exposure). The proportion of offspring per lineage retaining the same *agr* state as the initial founder cell, i.e., showing intergenerational stability of the *agr* state, is shown. Bars and dots represent mean and biological replicates, respectively. As only replicates with at least 8 lineages in a given founder *agr* state were included, displayed n varies. For the control, only in one replicate of *agr-IV* more than 8 lineages were in an activated *agr* state according to the founder cell classification and no activated founder cells for *agr-III* were detected in both AIP conditions. **B.** For the 64 nM homologous AIP exposure, the proportions of cells with switched *agr* state, defined by comparing the *agr* state at birth and division, is shown per generation (normalized to founder cell) and lineage, and separated by founder cell *agr* state. Dots are colored by the proportion of mother cells (the preceding generation) in activated *agr* state. For visualization purposes, random noise was added to the datapoints (20% and 1% jitter to the x- and y-axis values, respectively). The blue line represents a binomial regression which indicated no difference regarding founder cell state (p=0.26).

### Activation of the *agr* system increases bacterial generation time

QS activation typically coincides with low nutrient availability and resource diversion to virulence factor production. Therefore, we hypothesized that the physiological traits of *agr* activated and inactivated bacteria would differ. To explore this, we employed cell tracking to quantify cell size and generation time (timespan between divisions, Fig.3A) of bacteria exposed to 64 nM, a concentration which activated large proportions of the populations (Fig.1E). The generation times of bacteria not exposed to AIP displayed statistically significant differences among the *agr-*types with *agr-III* being the fastest and *agr-II* the slowest (median 22.00 and 35.75 min, Suppl. Fig.5A). 63% of *agr-II* bacteria exhibited an extended generation time (exceeding 60 min, GT60min+) during AIP exposure, which was significantly different from both the pre or post AIP phase (Fig.3B). In contrast, only a minority of bacteria from the other *agr-*types showed similarly high GT60min+, with *agr-III* bacteria barely having increased generation time (GT60min+ = 0.35%).

**Figure 5:**
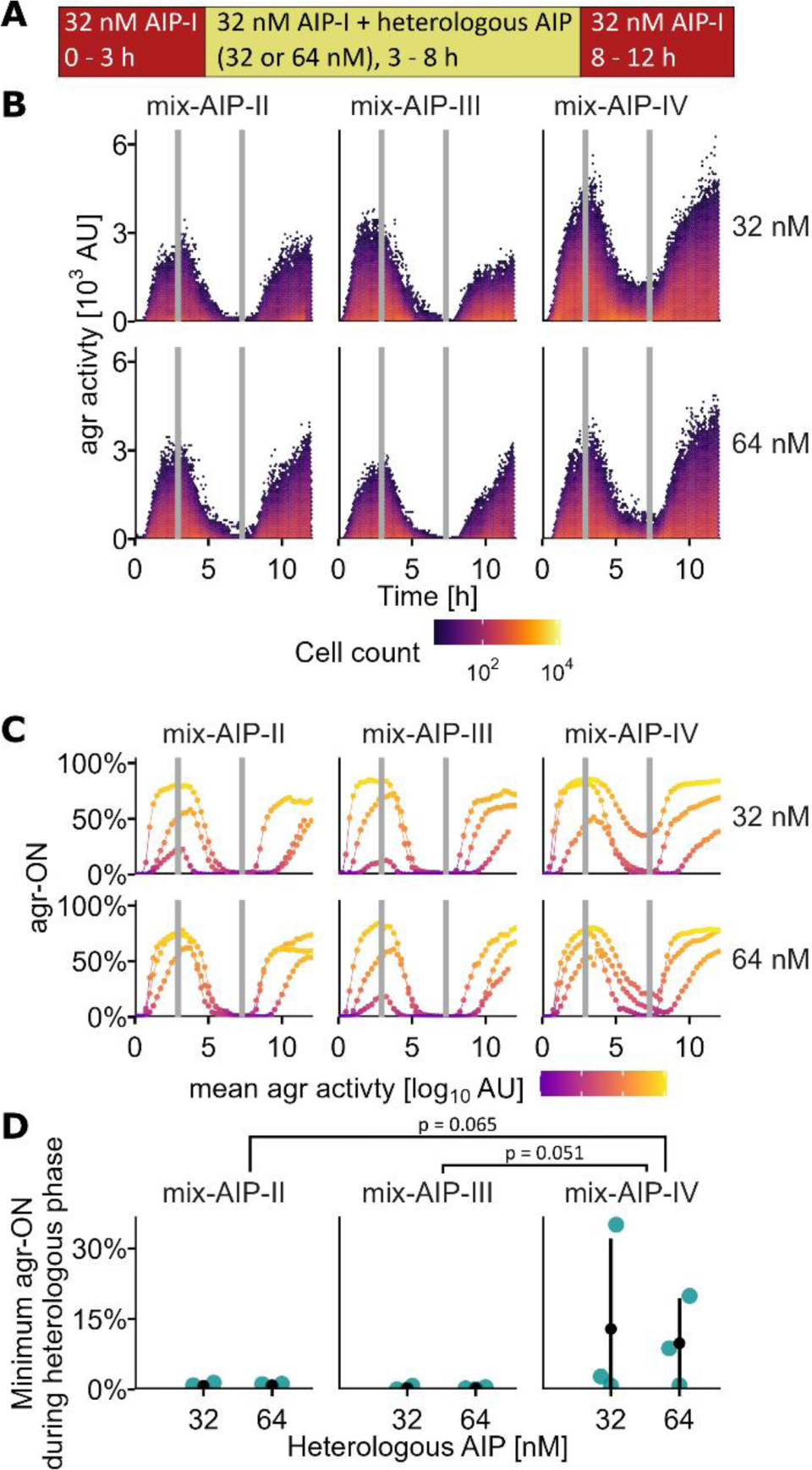
Heterologous AIP exposure efficiently suppresses *agr-I*. **A.** Using the mother-machine device, the *agr-I* strain underwent a sequence of AIP exposures. Experiments were performed in biological triplicates. **B.** *agr* activity of single cells (YFP fluorescence) during the three phases of AIP exposure. Switches are indicated by vertical gray lines. The type and concentration of heterologous AIPs in the combined exposure phase are labeled. Dot colors reflect log_10_ cell counts. On average, 203430 cells per condition were imaged (details in Suppl. Table2). **C.** Percentage of *agr* activated (ON) cells (YFP fluorescence > 35 AU) for each biological replicate, with exposure switches indicated by vertical gray lines. Color indicates average log_10_ *agr* activity. **D.** Lowest percentage of *agr* activated bacteria during the combined exposure phase. Blue dots represent biological replicates, black dots and bars show mean and standard deviation. Statistical significance was assessed with a Dunnett test using AIP-IV as reference.

Bacteria with extended generation times often had an activated *agr*, irrespective of their *agr*-type, evident when classifying bacteria by both these quantities (Fig.3C, compare upper quadrants). However, this was not mutually exclusive, as subpopulations of *agr* activated cells did not displayed extended generation times (Fig.3C, compare right-side quadrants).

The cell size increased significantly in bacteria with prolonged generation time, across all *agr*-types (Suppl. Fig.5B). A weak significant positive correlation (R^2^<0.065) between cell size and generation time was observed (Suppl. Fig 5C).

### Early AIP non-responders produce non-responder offspring

The observed *agr* bimodality (Fig.1E) could either be attributable to constant *agr* state switching of individual cells or to intergenerational stability of the *agr* state, acting as a form of non-genetic phenotypic inheritance^29,30^. Our experiments confirmed the latter. In the absence of AIP stimulation, offspring cells matched the inactivated *agr* state of lineage founder cells expectedly well (up to 100%), and intergenerational stability of the activated state was high during AIP stimulation (up to 96.7%, Fig.4A). *agr-IV* had the lowest intergenerational stability of the inactivated state (79.7%), potentially linked with the previously identified increased baseline *agr* activity without AIP stimulation (Fig.1D). The inactivated *agr* state in the presence of AIP stimulation was less stable between generations, indicating more frequent switches from inactivated to activated. Further, *agr* state switches mostly happened in early generations, regardless of the founder cell state, indicating rapid gene expression adaptation to the increased AIP concentration (Fig.4B). *agr-III* was again the exception and showed switches into the activated state late in the lineage, reflecting its low AIP sensitivity.

### Heterologous AIP suppress *agr-I* efficiently

Next, we utilized the mother-machine device and synthetic AIPs to study single-cell level *agr* interactions, focusing on *agr-I*. We observed that all tested heterologous AIP inhibited the *agr-I* system when it was first exposed to its own AIP (AIP-I) to pre-activate the *agr* system, then to a mix of AIP-I and a heterologous AIP, and finally back to AIP-I alone (Fig.5ABC, Suppl. Movies2). Inhibition occurred quickly, reaching the lowest proportion of activated bacteria (Fig.5D) within 2.2 to 4.4 h across all replicates. Reintroducing AIP-I alone promptly reactivated the *agr* system (Fig.5C). AIP-IV also inhibited the *agr-I* system, though its inhibitory strength, not significantly different from other heterologous AIPs, was generally weaker at both tested concentrations.

### Distinct *agr* interaction patterns across co-cultured bacterial communities

To study *agr* crosstalk of bacterial communities beyond pure AIP effects, we co-cultured the *agr-I* strain alongside each of the other *agr-*types in adjacent microfluidic chambers connected by nanochannels (Fig.6A). This design prevented mixing of cells across the nanochannels while allowing diffusion of soluble factors and accumulation of self-secreted AIPs leading to autoinduction of the *agr* system (Suppl. Fig.6). When changes in the *agr* activity of a community occurred, they often originated from the area nearest to the nanochannels and quickly spread throughout the entire chamber (representative kymographs in Suppl. Fig.7, Suppl. Movies3).

**Figure 6:**
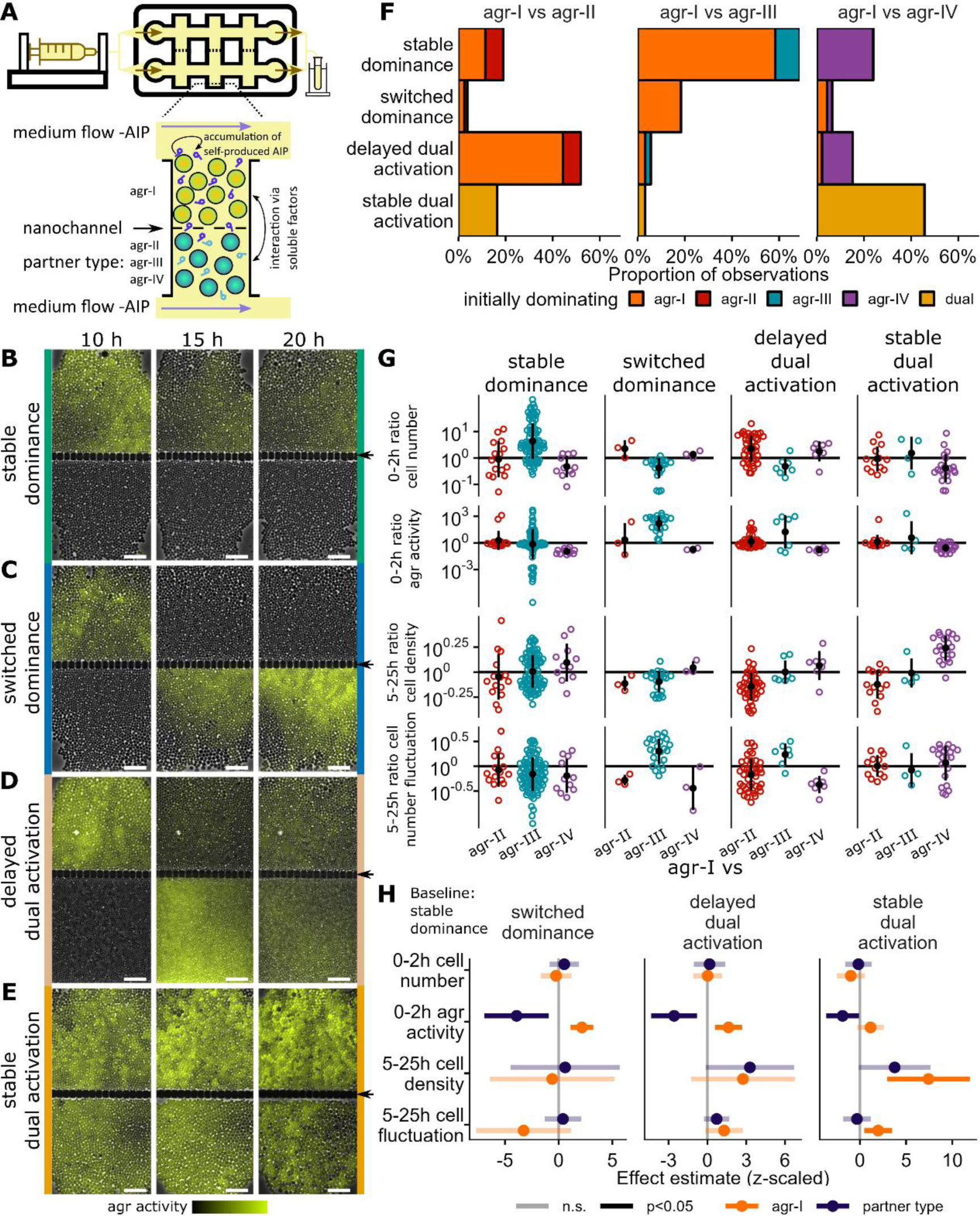
Four main classes of *agr* interaction dynamics in cross-talking populations. **A.** Schematic of connected-chamber microfluidic chip. Medium was independently provided at the top and bottom of chambers (size: 50×50 µm). The nanochannels separating the two chambers prevented mixing of the two populations while soluble factors produced by bacteria (including AIP) could diffuse through. *agr-I* was paired with either one of the other three types. Each pairing was repeated at least three times which included observation of up to 36 connected chambers, each sustaining bacterial communities of 1504 ± 420 cells. **B-E.** Representative images display the P3 *agr* activity (phase contrast overlayed with P3-YFP channel) at 10, 15, and 20 h post-inoculation for the four main categories of *agr* interaction dynamics. Arrows mark nanochannels and two different *agr* types are on each side of it. Scale bar: 10 µm. **B.** One population consistently displayed *agr* activation while the other remained suppressed. **C.** The dominating role switched between the two populations. **D.** Dual *agr* activation was seen with one population lagging or (**E.**) both populations maintaining a steady *agr* active state throughout the experiment. **F.** Distinct interaction categories emerged for each tested *agr-*type pairing. The *agr-*type which consistently or initially dominated is indicated by color. Number of observed chamber pairs of each class and *agr* combination can be seen in Suppl. Table3. **G.** Four population characteristics were quantified. Averages for the indicated timespans were calculated (Suppl. Fig.9) and ratio of *agr-I* to the partner *agr*-type is shown. One dot represents one connected chamber pair and is colored by partner *agr-*type. Black dots and bars show mean and standard deviation. **H.** Stable dominance, the most prevalent interaction observed across all *agr* combinations, served as the reference category in a multinomial regression analysis. All explanatory variables underwent z-scaling prior to analysis (raw values below and above the mean become negative and positive, respectively). Confidence intervals of the effect estimates are represented by bars, with the effect estimates illustrated as a circle, distinct for *agr-I* and the partner *agr-*type. Non-significant terms overlap with 0 and are indicated with transparent confidence intervals.

We categorized temporal *agr* interaction patterns into four main classes, observable across all *agr* combinations but with varying frequencies among the tested combinations. Three classes involved periods of an activated *agr* system of only one of the two interacting *agr-*types; either with no temporal change (’stable dominance’, Fig.6B, most common in *agr-I* vs *agr-III*, Fig.6F), temporal switching between the activated type (’switched dominance’, Fig.6C) or an initial phase of dominance followed by dual activation (’delayed dual activation’ Fig.6D, most common in *agr-I* vs *agr-II*, Fig.6F). The fourth pattern was characterized by constant dual activation (’stable dual activation’ Fig.6E, most common in *agr-I* vs *agr-IV*, Fig.6F). Two additional classes, ‘initial dominance’ and ‘unstable dual activation’, were only observable at low frequencies and in specific *agr* combinations (Suppl. Fig.8).

If phases of dominance occurred, the type which dominated differed between the tested *agr* combinations (Fig.6F). The *agr-I* dominated often against *agr-II* and *agr-III* but almost never against *agr-IV* with which dual activation was most common.

### Predictability of *agr* interaction categories by community characteristics

Four parameters describing the growth onset (0-2 h after inoculation) and maintenance (5-25 h after inoculation) phase of the bacterial populations were investigated regarding their predictability for the occurrence of the four interaction classes. 1) Onset cell number, potentially influencing total AIP production; 2) Onset *agr* activity, which might offer a ‘head start’ in AIP production due to the positive feedback loop of the *agr* system^31^; 3) Maintenance cell density, as higher densities could reduce diffusion and enhance AIP accumulation; 4) Maintenance cell number fluctuation, which could either influence AIP diffusion or *agr*-related cell dispersal due to biosurfactant production^32^ (Fig.6G).

In a multinomial regression, we compared interaction classes against ‘stable dominance’ as the baseline, finding significant associations with specific (z-scaled) parameters (Fig.6H). Including the two rare classes did not alter the qualitative model outcome but affected the significance of some parameters near the p=0.05 threshold (Suppl. Fig.10).

’Switched dominance’ was, compared to the baseline ‘stable dominance’, linked to high onset *agr* activity in *agr-I* and low onset *agr* activity in the partner type (Fig.6H), mostly due to the difference of *agr* activity in the combination with *agr-III* (Fig.6G). Onset *agr* activity was, compared to the baseline, also linked with ‘delayed dual activation’, which in addition was borderline significantly associated with both higher maintenance cell density of the partner type, especially in the combination with *agr-II* (Fig.6G), and higher maintenance cell number fluctuation of *agr-I*. ‘Stable dual activation’ showed similar associations but was additionally linked to increased late cell density of the *agr-I* type, markedly increased in the combination with *agr-IV* (Fig.6G).

## Discussion

Direct single-cell quantification of *S. aureus agr* dynamics within chemically controlled microfluidic environments, like those employed in our study, offer a significant advancement, providing novel insight into virulence regulation. Our findings revealed distinct AIP sensitivities across the four *agr*-types, with evident QS heterogeneity – unconfounded by AIP accumulation or spatial structure – likely explained by intergenerational phenotypic stability of the *agr* response. Intriguingly, *agr-I* was efficiently suppressed at the single-cell level by heterologous AIPs including the presumably cross-activating AIP-IV. Using novel microfluidic chip designs, our study also discerned four prevalent temporal *agr* interaction patterns within cross-talking neighboring populations, with their occurrence varying according to the specific interacting *agr* types.

In our analysis, *agr-III* emerged as the most AIP insensitive and *agr-II* as the most sensitive type. These findings offer nuanced insights adding to the inconclusive evidence on AIP sensitivity differences based on existing batch culture studies^14,33^. Our experiments suggest that the AIP concentration threshold for P3 activation is considerably higher for *agr-III* than for other types, implying *agr-III* activation only at high population density. We observed reduced baseline expression of the RNAII operon in *agr-III* and frequent switched dominance of it which both support this hypothesis and further indicate delayed AIP-III production. However, quantitative concentration measurement of such small peptides remains challenging with no established methodologies for AIP-III measurement to date^34,35^.

In addition to being the most sensitive *agr*-type, activated *agr-II* also showed increased generation time indicating an important physiological response, possibly reflecting a diversion of resources from growth to virulence factor procuction^36^ or cell wall damage^37^. High AIP concentrations typically indicate imminent nutrient scarcity due to increasing population density^38,39^. This premise creates a “signal confusion” for the bacteria in our experimental setting where both nutrient and AIP concentrations are high: while nutrient levels were favorable, high AIP concentration indirectly signaled potential nutrient depletion. We could argue that, in the case of *agr-II*, the nutrient limitation signal induced by *agr* activation supersedes actual nutrient availability, resulting in the observed extension of generation time. Our observations are further evidence for a complex interplay between metabolism and virulence regulation^39^.

We did not identify clear differential gene regulation of known regulators of the *agr* system among the four *agr*-types to explain AIP sensitivity differences and expect genetic background to be more influential. Nonetheless, two notable findings included the upregulation of *sarX* in *agr-II* and *IV*, a repressor of P2 and P3^6,24^ which would suggest low AIP sensitivity – the opposite of what we observed, and *sarT* downregulation in *agr-IV*, potentially explaining its slightly elevated baseline *agr* activity due to the negative feedback loop of *sarT* with *agr* via the P3 promoter^6,25^. The expression levels of *sarA* and *codY*, two key *agr* regulators^38,40^, were consistent across all *agr* types, suggesting other mechanisms at play affecting AIP sensitivity. Further studies are required to clarify the molecular mechanisms behind AIP sensitivity differences.

Our single-cell analysis underscores QS bimodality as an intrinsic feature of the *agr* system, independent of external factors such as AIP accumulation or metabolic alterations of the chemical properties of the environment. This addresses a critical gap left by earlier batch culture studies^16,17^, enhancing the understanding of QS heterogeneity. We also demonstrated that bimodality is most pronounced at intermediate AIP concentrations and is likely a consequence of intergenerational phenotypic stability of the *agr* activation state.

Contrary to results from bulk studies traditionally using cell-free spent growth medium^13^, AIP-IV exhibited a significant yet incomplete inhibitory effect on *agr-I*, highlighting the sensitivity of single-cell assays and the importance of using pure AIPs in detecting subtle QS interactions. Our results thus emphasize that inhibition by heterologous AIP is less influenced by structural differences than activation^20,33,41^, which may be even more sensitive to changes than previously anticipated.

We identified four common *agr* interaction patterns across all tested *agr*-type combinations with their occurrence rates varying by *agr* combination. Stable *agr* dominance was common, especially *agr-I* dominating against *agr-III*, aligning with earlier observations^14^, and our own data indicated rapid initial activation of *agr-I*. However, the frequent occurrence of switched dominance in the same pairing suggests that once activated, *agr-III* maintains its *agr* activity more robustly than *agr-I* and can even suppress it. Dual activation was notably frequent between *agr-I* and *agr-IV*, corresponding with previous reports of cross-activation of bulk cultures^13^ or the partial inhibition demonstrated in our study. The early *agr* activity during the population establishment phase significantly influenced the interaction dynamics, underscoring the critical role of initial gene expression in processes such as niche colonization or barrier breaches^42,43^. The difficulty in predicting interaction outcomes illustrates the complex and often stochastic behavior within bacterial communities.

Our study advances the understanding of *S. aureus* QS dynamics through innovative microfluidic experiments yet emphasizes the necessity for broader investigation across diverse strains to confirm these insights regarding AIP sensitivity and *agr* interaction patterns beyond specific genetic backgrounds. The mother-machine platform and connected chamber devices, coupled with our image-analysis pipeline, sets the stage for exploring QS inhibitor efficacy^44,45^ and dissecting the role of commensal bacteria in QS regulation. Highlighting the significance of distinct AIP sensitivities and *agr* interaction patterns, our findings advocate for targeted, personalized approaches in treatment strategies that consider the virulence and genetic diversity of *S. aureus* strains^46^.

Our research indicates that treatments aiming to suppress virulence, particularly focusing on *agr-II* due to its heightened AIP sensitivity, could improve infection management and indirectly minimize antibiotic usage. However, clinical data correlating *agr*-type with infection severity remains elusive and is possibly influenced by regional factors^47–49^ and the genetic background of *S. aureus* strains^12^. Pursuing further studies with clinical isolates to correlate *agr* AIP sensitivity with infection outcomes could validate the clinical relevance of our discoveries. By providing a more detailed understanding of QS dynamics, our work contributes to a strategic framework for inhibiting the shift of *S. aureus* from a benign colonizer to a threatening pathogen, aligning with the broader goal of devising more effective and sustainable approaches to bacterial infection control.

## Methods

### Strains and growth conditions

We used four distinct *agr* prototype *S. aureus* strains^7^ (LAC: *agr-I*, ST8; 502a: *agr-II*, ST5; MW2: *agr-III*, ST1; MNTG: *agr-IV*, ST2276) containing the pDB59^50^ *agr* P3-YFP reporter plasmid, provided by Alexander Horswill. For overnight cultures, single colonies were inoculated into 3 ml Tryptic Soy Broth (TSB, BD) with 10 µg/ml chloramphenicol (Cm10) for plasmid maintenance and 0.01% Tween20 to minimize aggregate and biofilm formation and incubated at 37 °C with 220 rpm shaking. We purchased synthetic AIPs from Cambridge Research Biochemicals, stored initial DMSO stocks at -20° C, and diluted them in Phosphate Buffered Saline (PBS) before addition into the growth medium at specified concentrations.

### Mother-machine microfluidic device

Molds of the microfluidic device were provided by Martin Ackermann, with device design and fabrication methods described elsewhere^51^. In brief, the design features six independent replicates of hundreds of chambers (height: 0.93 µm, length: 25 µm, width: varying from 1.2 to 1.6 µm), arranged perpendicular to a main flow channel (height: 21 µm width: 200 µm). For device fabrication, polydimethylsiloxane (PDMS, Sylgard 184 Silicone Elastomer Kit, Dow Corning) was mixed at a 1:10 ratio with its curing agent, poured on the mold, degassed for 10 min, and cured at 80° C for one hour. Following, the PDMS device was cut out, and inlet and outlet holes (0.75 mm diameter) were created using biopsy punchers (WellTech). The device was then bonded to a #1.5 glass slide (Corning) via plasma-treatment.

### Connected chambers microfluidic device

The microfluidic device molds were created through lithography of SU8 (Microchem, Germany) on silicon wafers. The chip design incorporates two primary channels for individual strain and media injections (height:20 µm, width: 100 µm), linked by a grid of 100 chamber pairs (height: 0.85 µm, length and width: 50 µm). Barrier nanochannels (width: 0.4 µm, length: 3 µm, spaced 2 µm apart) separate each chamber pair, allowing solute transfer but preventing cell translocation. This design facilitates co-culturing of different strains in adjacent chambers, with hundreds of replicates per chip. Additionally, isolated dead-end chambers act as controls. For device fabrication, PDMS was mixed at a 1:5 ratio with its curing agent, poured on the mold, degassed for 20 min, and cured at 80° C for two hours. After cutting, the PDMS device was heated at 120° C for 20 min. Inlet and outlet holes (1.5 mm diameter) were punched, followed by an isopropanol wash, and the device was plasma-bonded to a #1.5 glass slide (VWR).

### Microfluidic device inoculation

Overnight cultures were diluted 1:50 in fresh 4 ml of the same medium and re-incubated for 2 h. Afterward, they were washed twice with TSB containing 0.1% Tween20 and then concentrated into 80 µl to achieve a dense bacterial suspension. The suspension was loaded into the microfluidic device from the outlet side with a pipette. The outlet and inlets were connected to a waste collection and a syringe pump (KF Technology, NE 1600, 10 ml syringes), respectively. We used PTFE (0.3 mm inner, 0.76 mm outer diameter, adtech) tubing for mother-machines and medical grade non-DEHP polymer (0.51 mm inner, 1.52 mm outer diameter, Tygon) for connected chamber chips.

For the mother-machine experiments, TSB with Cm10 and 0.01% Tween20 was used, flowing at 0.5 ml/h. After loading, cells were left overnight to ensure metabolic recovery and *agr* synchronization before experimental AIP exposure. In contrast, we began immediate imaging of the connected chamber chips to capture community development from their initiation. For these chips, we used a 1:1 mix of TSB and PBS, with Cm10 and 0.01% Tween20, at a 0.25 ml/h flow rate. This lower nutrient and flow environment aimed to reduce cell division rates and minimize AIP washout. All experiments were performed at 37 °C, with medium changes executed by manual syringe swaps at specified timepoints.

### Image acquisition

We used a fully automated Olympus IX83 P2ZF inverted microscope, controlled via CellSense 3.2 (Olympus), for imaging. Phase contrast (200 ms exposure, 130 intensity IX3 LED) and YFP fluorescence (200 ms exposure, 20% intensity 500 nm CoolLED pE-4000, triple-band filter for CFP/YFP/mCherry) images were captured at 16-bit resolution in intervals ranging from 2 to 15 min using a 100×, 1.3 numerical aperture oil phase objective (Olympus) and an ORCA-flash 4.0 sCMOS camera (Hamamatsu). Up to 16 positions for each of the six replicates of mother-machine devices per chip and up to 36 positions per replicate on connected barrier chips were manually selected.

### Image analysis

We developed an image analysis pipeline operating solely on phase contrast images for improved generalizability, which also allows all fluorescence channels to be used for reporter quantification (here, YFP for *agr*-P3 quantification). Initial steps were performed in MATLAB (MathWorks) and included image registration and chamber detection (detailed description in Suppl. Methods and Suppl. Fig.11). Subsequent steps involved single-cell segmentation using StarDist (version 0.8.3)^52^ and tracking with a modified version of DeLTA (version 2.0)^53^, both executed in Python (versions 3.7 and 3.9, respectively).

During our initial assessment of the overview movies produced in MATLAB, we manually discarded positions affected by image registration errors, chamber detection issues, high rates of out-of-focus frames, or obvious cell death, potentially attributable to media switch-induced pressure changes.

Given the suboptimal performance of screened pre-trained StarDist models, we derived two specific segmentation models for the distinct microfluidic device designs. The differences in cell density and image complexity between the mother-machine and connected chamber chips necessitated this approach. We generated 707 and 508 new segmentation training images for mother-machine (one image containing one chamber) and connected chamber chips (random 100×100 pixel crops), respectively (Suppl. Fig.12A-D). Initially, a set of 100 images was manually annotated using LABKIT^54^ in FIJI-imageJ and a preliminary StarDist model trained on these. This model was applied to batches of 100 images, undergoing manual corrections before a new model was trained from scratch with the complete set of images. For re-training, we used the following parameters: 64 rays, 64×64 pixel patches, validation split ratio 0.125, batch size 8, 400 epochs, 100 steps per epoch. Data augmentation included random flips and rotations as well as random intensity changes and addition of noise. Only the best model regarding validation performance (loss) was saved. The achieved segmentation precision was satisfactory for both chamber designs (Suppl. Fig 12EF). Cell properties including fluorescence channel quantifications were measured with regionprops of scikit-image^55^. For mother-machine experiments, segmented cells were further filtered according to cell size to include only cells with area between 0.3 and 2.5 µm^2^ and interpolated volume between 0.1 and 3 µm^3^. For response time calculation, only replicates reaching the threshold of 50% activated bacteria were included. For connected chamber experiments, only chambers with a median cell number of 700 or higher were included.

Due to the high cell density in barrier-chips, cell tracking was at the time of writing impossible without extremely high image acquisition rates which we omitted. Most bacterial cell tracking pipelines, including DeLTA 2.0 itself, were tailored for rod-shaped bacteria and the pretrained model did not perform well with cocci. Performing iterative rounds of prediction, correction and training, we created 11’671 new tracking event training sets specifically for cocci in mother-machine devices with scripts provided by Owen M. O’Connor^53^. We additionally altered the original weights function to a more uniform distribution. Upon combining our new training data with 4’732 randomly selected sets from the original publication, we re-trained the deep-learning tracking model with the following parameters: validation split ratio 0.125, batch size 8, 900 epochs, steps per epoch 1500 (200 for validation) and early stopping patience of 30 epochs. Default data augmentation of DeLTA 2.0 was used and only the best model regarding validation performance (loss) was saved.

We modified DeLTA 2.0 to use StarDist2D segmentations instead of the built in segmentation algorithm due to increased segmentation accuracy. The obtained output of applying the tracking model, including lineage information and fluorescence intensity values, was subsequently transformed into Python pandas tables, leveraging code provided by Simon van Vliet^56^. For our analysis, we only included cells for which a mother cell and a division event were observed and cells with a tracked lifespan below 10min or above 6h were excluded. For assessment of intergenerational stability, only replicates with at least 8 lineages in a given founder *agr* state were included.

Our data presentation is as follows: Fig.1, 5, and 6 display StarDist single-cell segmentation data. While Fig.1D and 5B depict the raw single-cell segmentation data, other panels present aggregated data as per indicated parameters. Fig.3 and 4 feature DeLTA 2.0 tracking data. Single-cell data from connected chamber chips was aggregated into descriptive statistics, either for 6.5×6.5 µm bins or for the entire chamber. We focused on the timespan of 5 to 25 h post-inoculation of the experiment to classify *agr* interaction patterns. Communities were classified into *’agr* ON’ and *’agr* OFF’ based on average fluorescence intensity per channel. Detailed description of the categorization scheme of the dynamics can be found in the Suppl. Methods.

### Whole-genome sequencing and comparative genomics

DNA was isolated from bacterial colonies grown on TSB with Cm10 and libraries prepared with the QIASeq FX DNA library kit (Qiagen) and sequenced with a MiSeq instrument using MiSeq reagent kit V2 (Illumina). 150 bp paired-end reads were assembled with SPAdes (v3.15.5^57^) with the careful argument and default k-mer size. Annotation was performed with prokka (v1.14.6^58^), and gene presence-absence analyzed with Roary (v3.13.0^59^) using https://github.com/judithbergada/Pipeline_GenomeAnalysis by Judith Bergadà Pijuan.

### RNA sequencing and analysis

To assess baseline gene expression profiles of all four strains, overnight cultures grown at 37° C in TSB with Cm10 and 0.01% Tween20 were diluted 1:20 in fresh medium and regrown for 1.5 h. Bacteria were washed twice in RNAlater and RNA was isolated using the RNeasy Mini kit (Qiagen) following manufacturer’s protocol including DNAse treatment (Turbo DNase kit, Invitrogen). Paired end 150 bp RNAseq was performed without rRNA depletion by the Functional Genomics Center Zurich (FGCZ) of University of Zurich and ETH Zurich using the TruSeq RNA Library Prep Kit on an Illumina NovaSeq X Plus yielding on average 47.6 million reads per sample. Analysis was performed using a modified prokseq pipeline (v2.0^60^), using default parameters. In short, quality control and filtering were performed using AfterQC (default parameters, v0.9.7^61^), followed by alignment to the reference *S. aureus* USA300 FPR3757 genome (GenBank <u>CP000255.1</u>) with bowtie2 (v2.3.5.1^62^) and subsequent execution of featureCounts (subread v1.4.6^63^). These RNA read counts were imported in R and analyzed using DESeq2 (v1.42.0^64^) and apeglm^65^. Gene annotation of the reference genome was enhanced using the pan genome annotation of AureoWiki^66^.

### Statistical analysis

Statistical analysis and figure creation was performed with R (v4.3.0), R Studio and tidyverse^67^. Response time to AIP concentration changes of independent biological replicates was assessed with linear regression using the interaction of *agr-*type and log2 AIP concentration followed by estimated marginal means (emm)^68^ *post-hoc* tests using multivariate *t* distribution (mvt) based *p*-value correction (Fig.1FG). Generation time of individual bacteria was log10 transformed and assessed with a mixed effect model using a random intercept term for biological replicate and the three-way interaction of *agr-*type, AIP concentration and experimental phase, followed by emm *post-hoc* tests using mvt correction (Fig.3B). Cell area was assessed with the same model structure but using the two-way interaction of AIP concentration and generation time cutoff (Suppl. Fig.5B). The correlation between generation time and cell area of individual bacteria (Suppl. Fig.5C) was quantified with Kendall correlations. The proportion of switched *agr* state of bacterial lineages was analyzed with a binomial regression using the three parameters normalized generation, *agr-*type, and founder cell state, and the effect of the founder cell parameter is reported (Fig.4B). Differential expression (RNAseq) of three independent exponential cultures was analyzed with DESeq2 using default parameters (Fig.2, Suppl. Fig.4). The minimum proportion of activated bacteria among independent replicates was assessed with a Dunnett test (Fig.5D). Connected chamber chip dynamics categories were assessed with multinomial regression^69^ using the four described parameters (Fig.6H, Suppl. Fig.10). Details of the statistical analysis can be found in Suppl. Data1 and Suppl. Data2.

### Data availability

Example movies of one position for each experimental condition can be found in the supplemental material. Raw images of all example movies including the output of the fully executed bioimage analysis pipeline are on BioImage Archive with the accession number S-BIAD1046 (https://www.ebi.ac.uk/biostudies/bioimages/studies/S-BIAD1046), additional raw images or processed data are available upon request. StarDist2D segmentation models for the mother-machine and connected chamber chips as well as the retrained DeLTA 2.0 tracking model and the used training data for these three models can be found on Zenodo (https://zenodo.org/doi/10.5281/zenodo.10694024). DNA and RNA sequencing data are deposited on the public archives European Nucleotide Archive (ENA) at EMBL-EBI under accession number PRJEB72799 (https://www.ebi.ac.uk/ena/browser/view/PRJEB72799) and the ArrayExpress database under accession number E-MTAB-13859 (https://www.ebi.ac.uk/biostudies/arrayexpress/studies/E-MTAB-13859), respectively.

### Code availability

Code for image preprocessing (registration, area of interest detection, crop and copy single-chamber images, overview movie generation) as well as the code for automated batch cell segmentation and cell tracking are deposited on GitHub (https://github.com/JulBaer/azimages, release v1.0.0).

## Supporting information

Supplemental Material (Figures, Tables, Movie and Animation Legends, Methods)

Supplemental Animation 1

## Acknowledgments

We thank Alexander Horswill for providing the *agr* prototype strains including the pDB59 plasmid, Martin Ackermann for providing the mold of the mother-machine, Clément Vulin for initial help establishing mother-machine experiments, Yohanes Belay Mignote for assistance in generating segmentation training images, Owen O’Connor for his help in modifying the original DeLTA 2.0 code to integrate external segmentations, Simon van Vliet for providing code to transform DeLTA 2.0 output into Python pandas tables, and Federica Andreoni and Mariane Pivard for proof-reading.

## Funding

A.S.Z. was supported by the Swiss National Science Foundation (SNSF) project grant 310030_204343, E.S. by the SNSF PRIMA grant 179834, G.S.U. by the NCCR Microbiomes (SNSF) grant 180575 and S.G.V.C. by the Marie Skłodowska-Curie (MSCA) individual fellowship grant 101033169.

## Declaration of Generative AI and AI-assisted technologies in the writing process

The manuscript was written by the authors. ChatGPT (GPT-4, last accessed at 16. February 2024) by OpenAI was employed for grammar correction, text streamlining and improving language clarity, and as a tool to explore alternative phrasings. It is important to note that ChatGPT was not used for analysis, intellectual content creation or literature review. The authors have thoroughly reviewed, verified and edited any passages generated by ChatGPT, taking full responsibility for the manuscript and overall quality and accuracy.

## Notes

### Competing Interest Statement

The authors have declared no competing interest.

