## Supplemental Material (Figures, Tables, Movie and Animation Legends, Methods) for "Single-cell approach dissecting *agr* quorum sensing dynamics in *Staphylococcus aureus*"

### Supplemental Figures

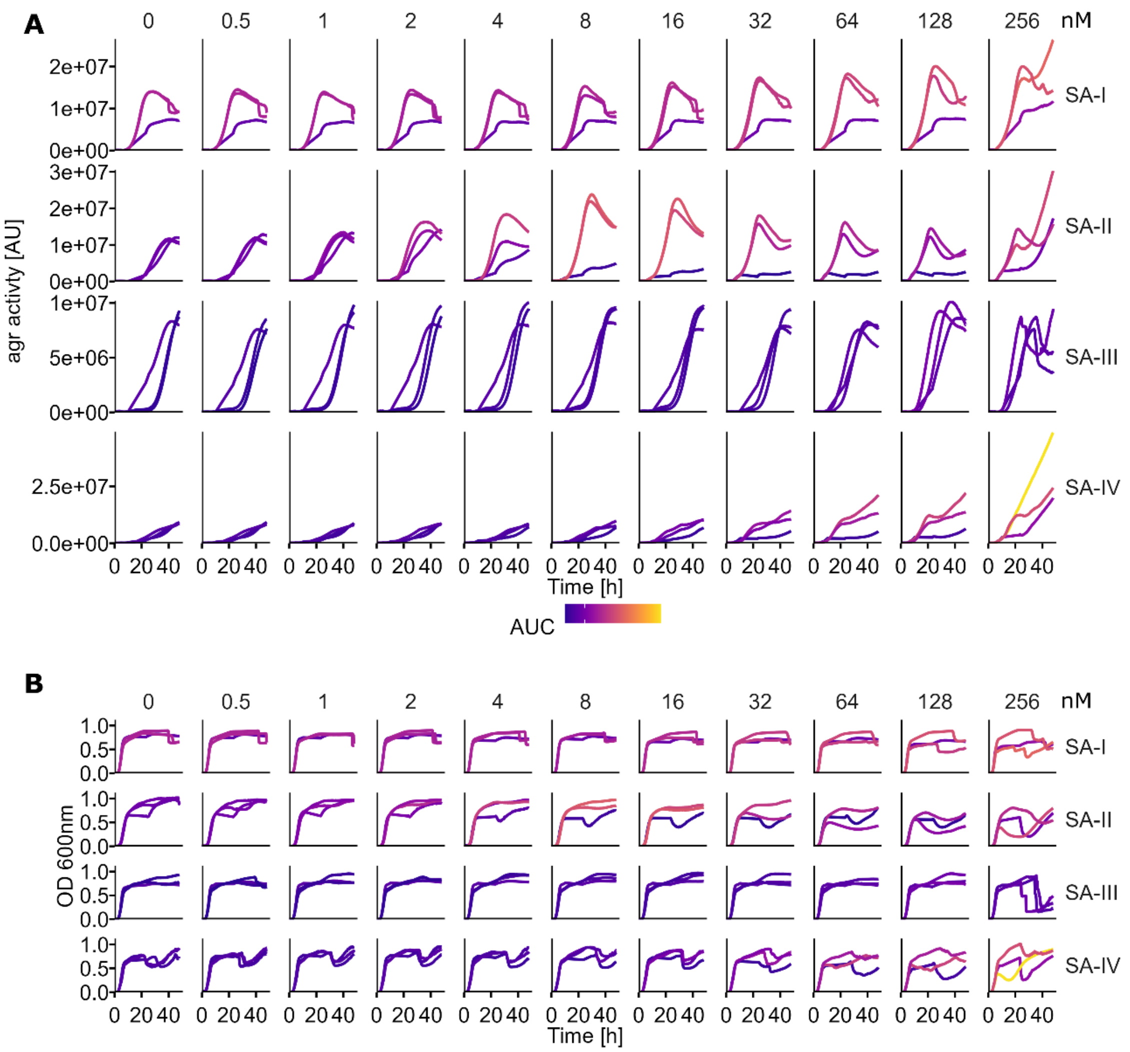


**Supplemental Figure 1: Plate-reader based *agr* activity of the four *agr*-types. A.** *agr* activity was measured over 48 h in a plate-reader with increasing concentrations of AIP. Increased fluorescence and earlier onset were observed at high concentrations. Area under the curve (AUC) was calculated per replicate and used to color-code the curves**. B.** Optical density was measured simultaneously. The same color-code as in A was used. High concentrations seemed to reduce the OD and were likely attributed to reduced attachment on the well bottom.

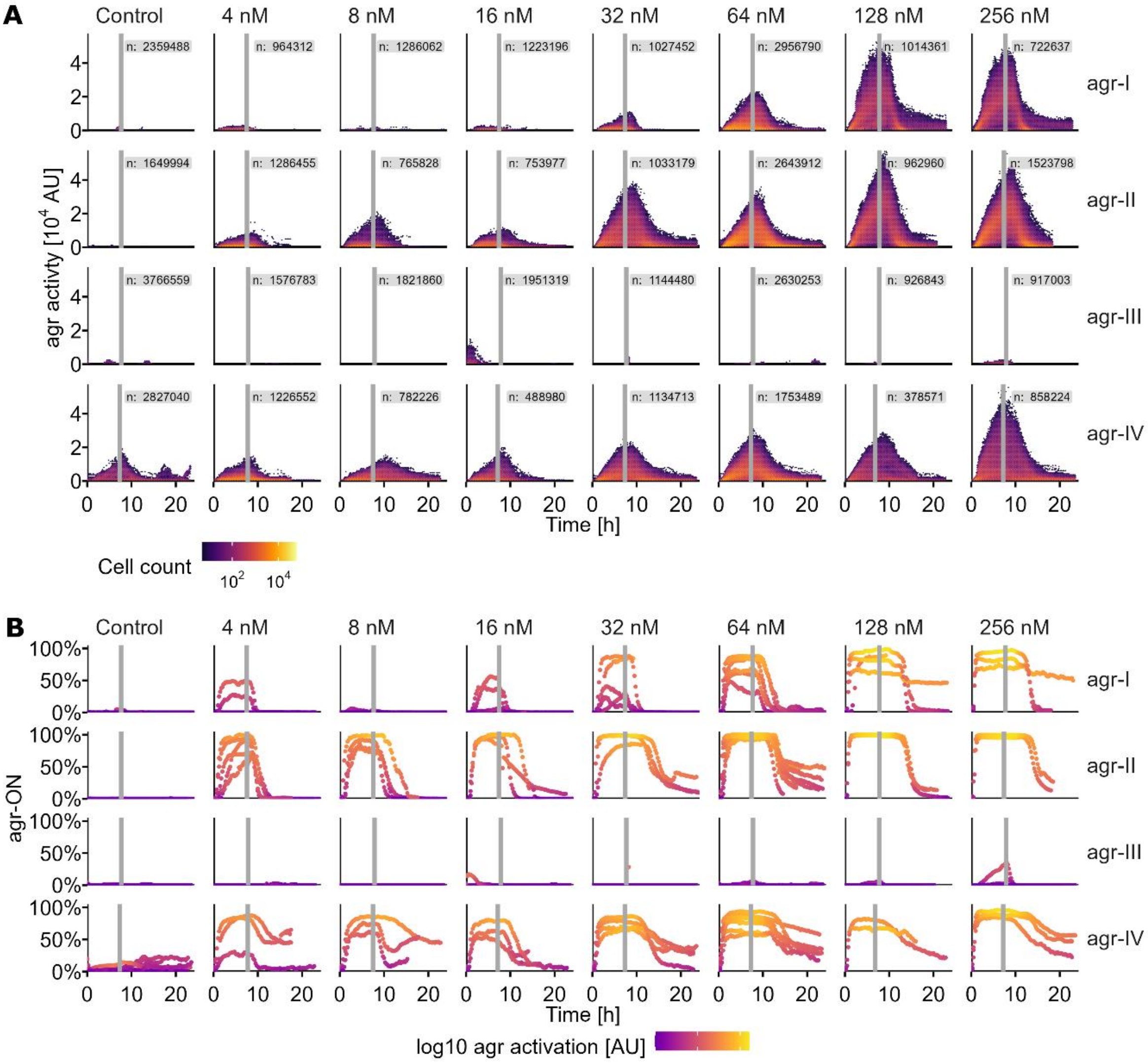


**Supplemental Figure 2: Additional homologous AIP concentrations. A.** Single-cell *agr* activity during AIP exposure (concentration annotated above each column) and after AIP removal (indicated by gray lines). Each row represents one *agr*-type. Dot color reflects $\log_{10}$ cell counts, n equals total counts per panel labeled. Control, 16 and 64 nM are already shown in Fig. 1D and are repeated here for completeness. **B.** Proportion of *agr*-ON cells (YFP signal per cell > 35 AU) calculated per biological replicate. AIP removal is indicated with a vertical gray line and exogenous AIP concentration annotated at top of panels. As SA *agr-III* cells respond very weakly to exogenous AIP, the threshold of 35 AU is almost never reached.

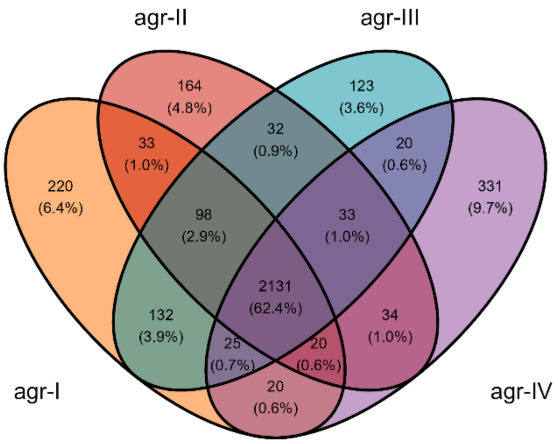


**Supplemental Figure 3: Venn diagram of gene presence-absence data of the four *agr*-types.** Whole genome sequencing Venn diagram based on annotation with Prokka and Roary.

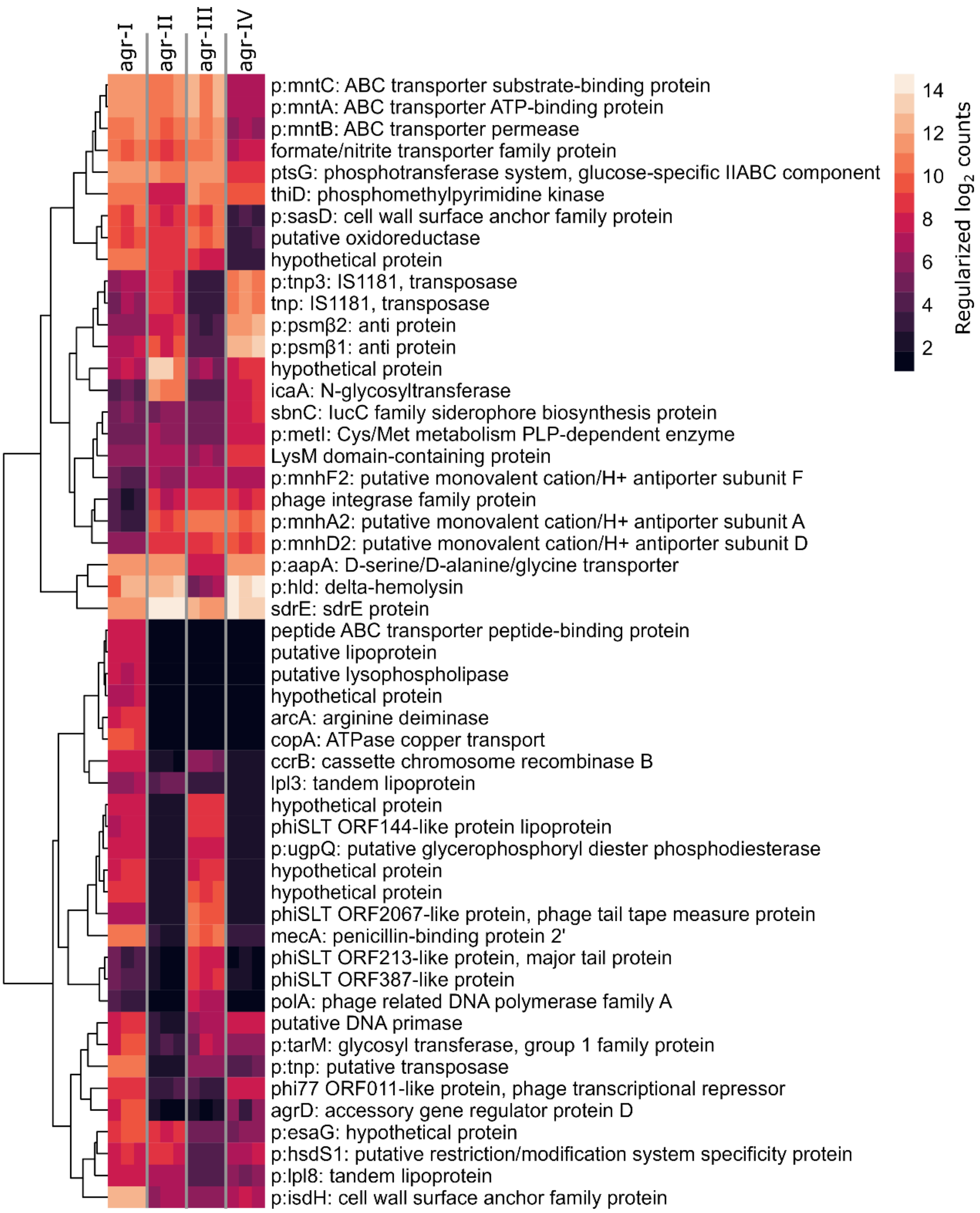


**Supplemental Figure 4: Identified differentially expressed genes.** RNA sequencing of exponentially growing cultures without AIP stimulation was performed to quantify baseline gene expression profiles. Heatmap of regularized log transformed RNA counts of the 25 most significant genes for each pairwise comparison of *agr-I* with any of the other three *agr*-types. Columns represent biological replicates. Gene names starting with “p:” are originating from the pan genome annotation of aureowiki. Genes without a gene name did not have an annotation in the genome of USA300_FPR3757 on aureowiki.

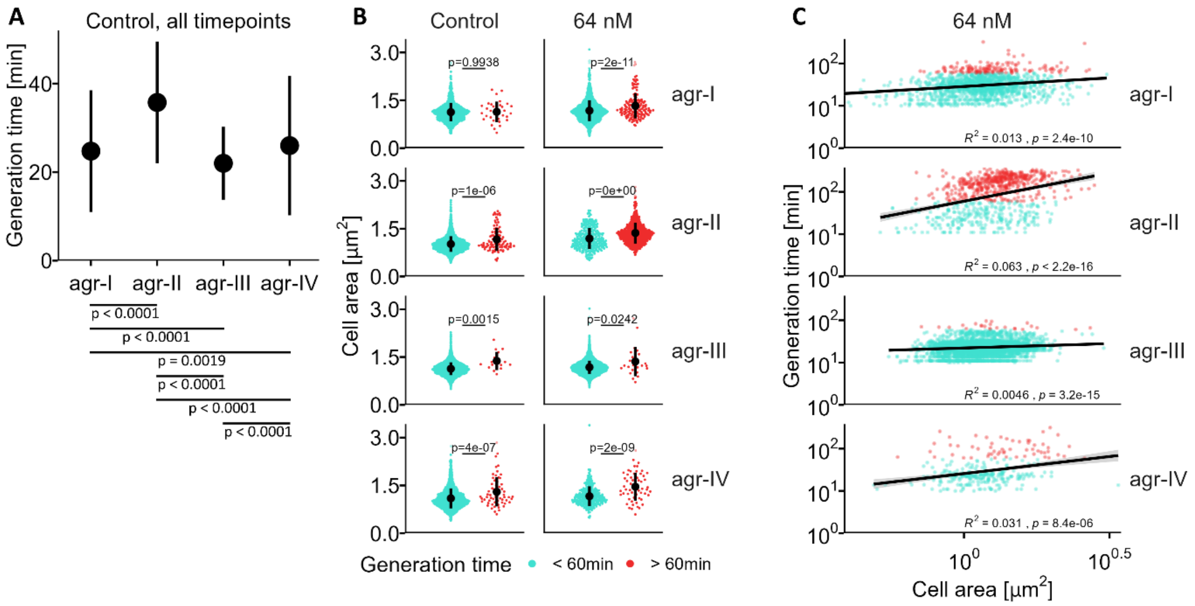


**Supplemental Figure 5: Cell area for normally and slowly dividing cells. A.** Median ± interquartile range calculated by combining single cell data of all three timepoints of the control experiments shown in Fig. 3B. Statistical significance was assessed with a mixed effects model on $\log_{10}$ transformed generation time followed by estimated marginal means pairwise comparisons (N cells analyzed for the four *agr*-types: 2662, 5093, 12957 and 3443). **B.** Maximal cell area during AIP exposure, separated by the generation time threshold of 60 min. Black dot and bar show mean and standard deviation. **C.** Scatterplot of generation time versus cell area (both $\log_{10}$ transformed), colored by generation time threshold of 60 min. Kendall correlation p-value and $R^2$ is annotated.

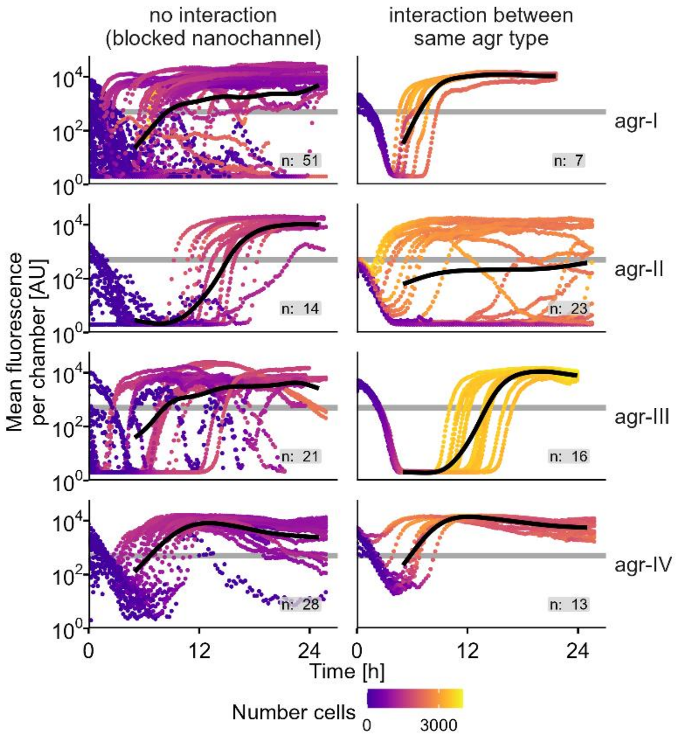


**Supplemental Figure 6: Control experiments for the connected chamber chips.** Left column shows data from chambers without the possibility of interaction via diffusion of metabolites (control chambers at opposite ends of flow channels or with a solid wall instead of permeable nanochannels). Right column data is from regular nanochannel chamber experiments with the same *agr*-type on both sides. Number of chambers is annotated.

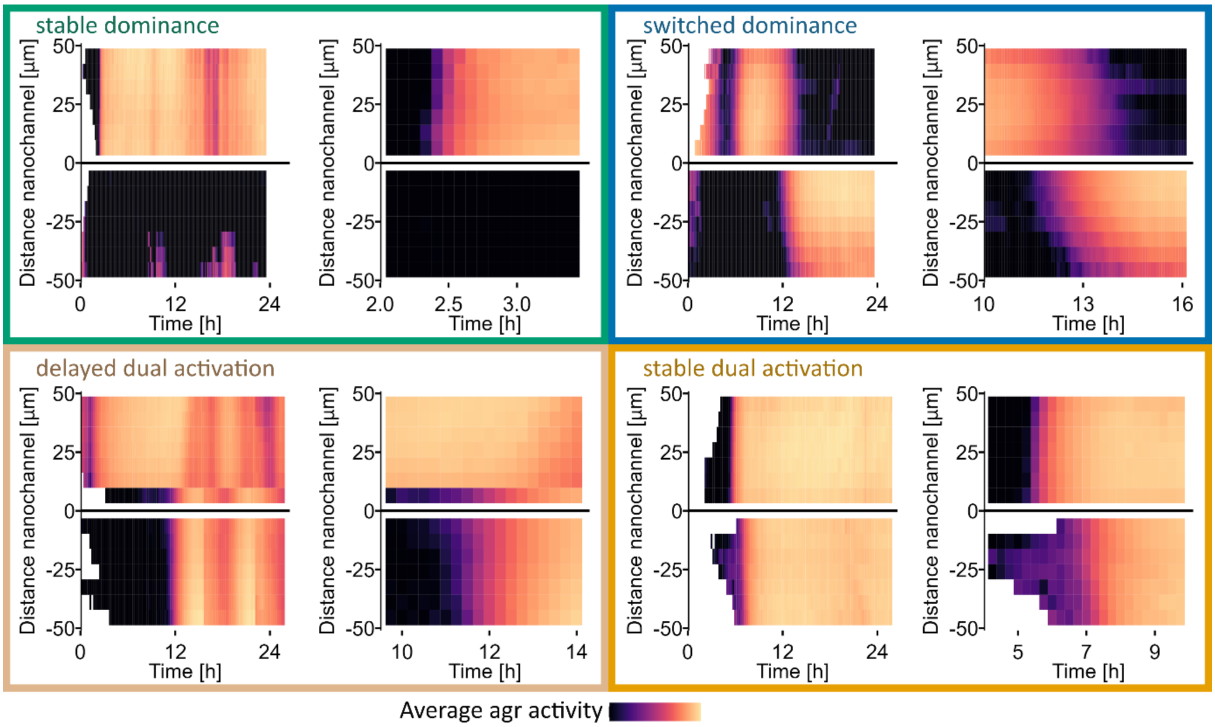


**Supplemental Figure 7: Representative kymographs of the four most frequent categories of *agr* interaction dynamics** (same examples as shown in Fig. 6B-E). Color intensity represents average *agr* activity as a function of time and distance from the nanochannels. White sections indicate areas with low number of cells. For each category, the left kymograph shows the full duration of the experiment, and the right is a zoom showing the timeframe at which changes in overall *agr* activity occurred. The *agr-I* strain is always on top (positive distance from nanochannel) and the partner *agr*-type at the bottom (negative distance from nanochannel).

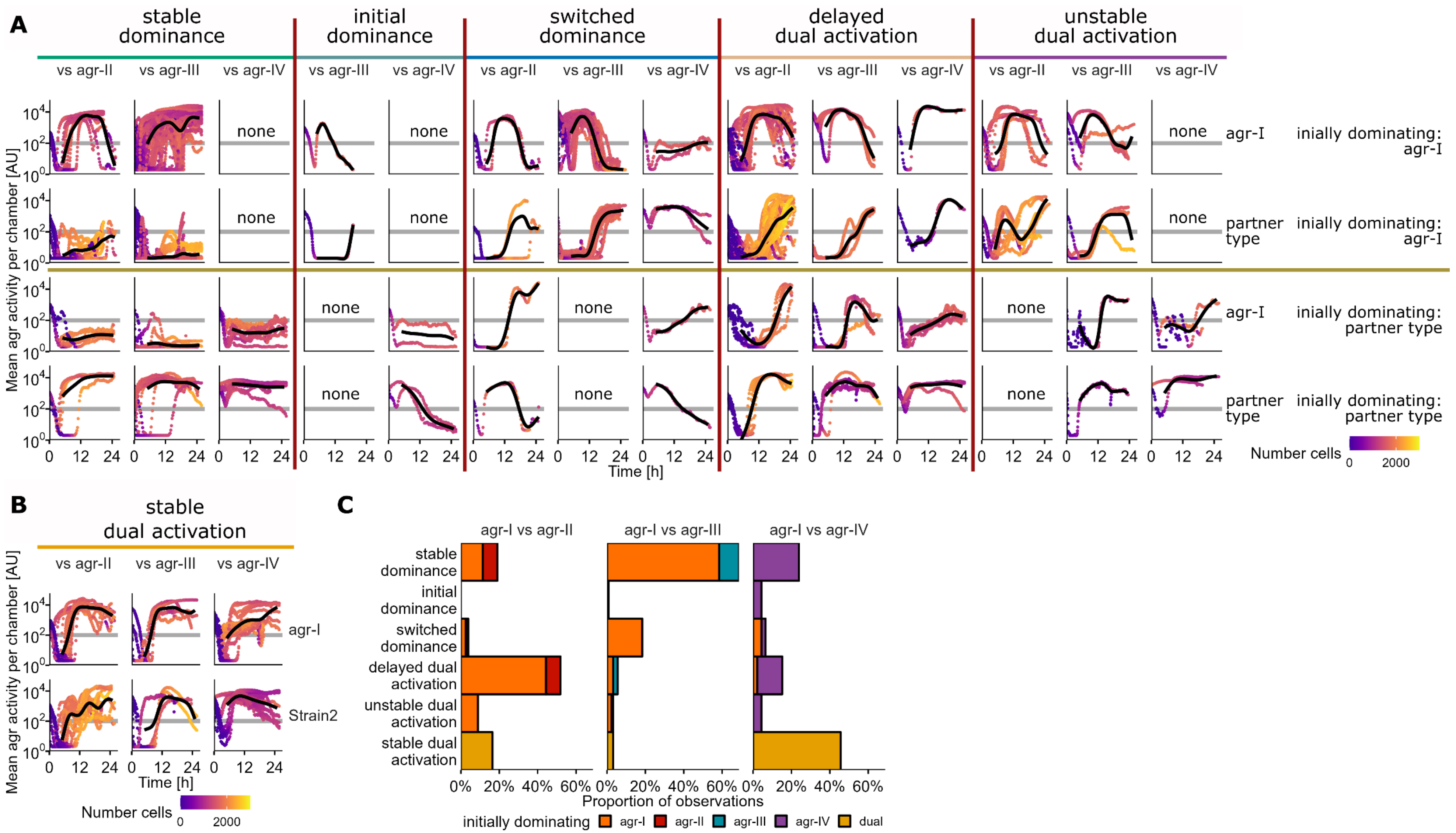


**Supplemental Figure 8: Complete set of all interaction chamber data. A.** Mean *agr* activity of all observed interactions. Each two rows belong to the same nanochannel connected chamber *agr* combination (divided by brown horizontal lines). The top of these two is always showing data of *agr-I* and the bottom row shows data of the partner *agr*-type annotated above each row in the vs statement. The top two rows are showing data in which *agr-I* was the (initially) dominating strain. Columns indicate the dynamic category (divided by red vertical lines) and *agr* combination. **B.** Same layout as in A for the stable dual activation category which lacks a dominating *agr*-type. **C.** Distinct interaction categories emerged for each tested *agr*-type pairing. The *agr*-type which consistently or initially dominated is indicated by color. Parts of panel C are shown in Fig. 6F and are repeated here for completeness.

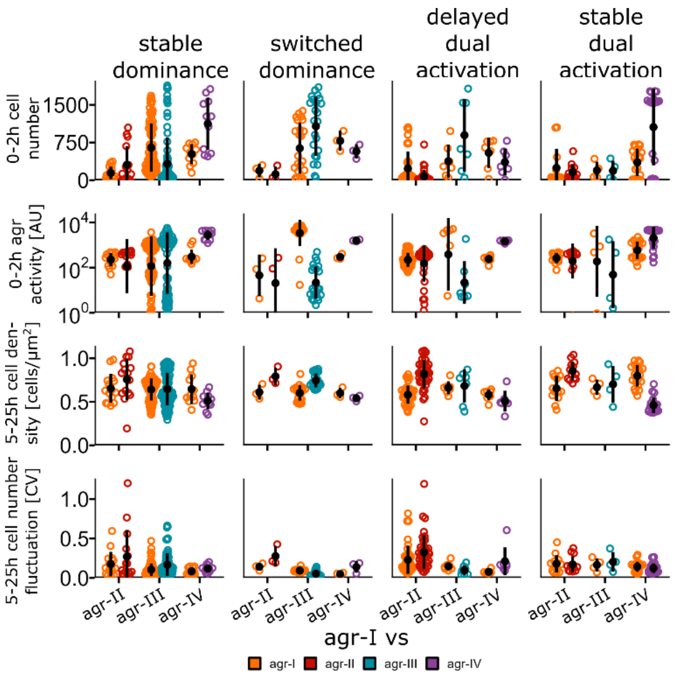


**Supplemental Figure 9: Raw data of the four parameters used in the multinomial regression.** This shows the same data as Fig.6G. Averages for the indicated timespans are shown. One dot represents one connected chamber pair and is colored by partner *agr*-type. Black dots and bars show mean and standard deviation.

CV = coefficient of variation.

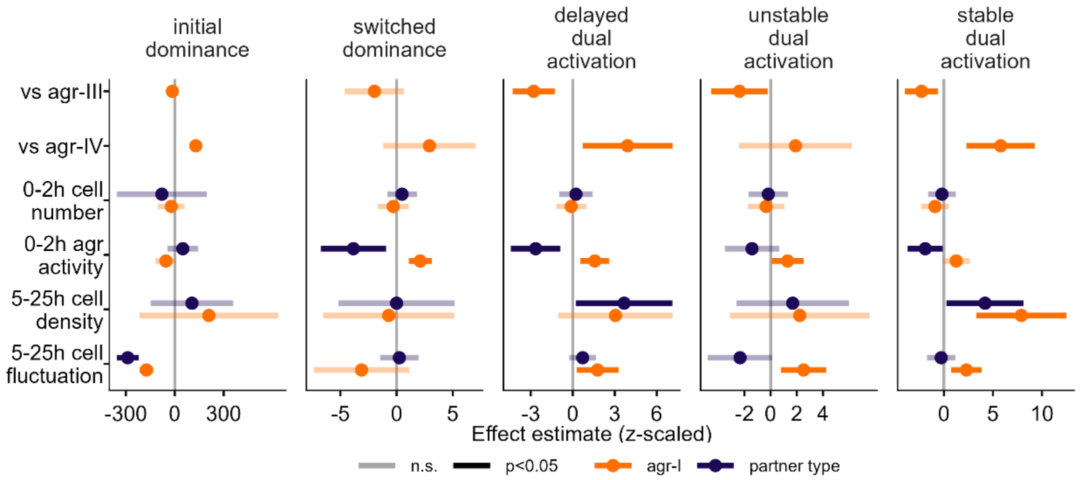


**Supplemental Figure 10: Complete multinomial regression analysis of the dynamic categories.** This shows the same model type as in Fig. 6H but including the two rare interaction classes and additionally visualizing the effect of *agr*-type combination. Stable dominance, the most prevalent interaction observed across all *agr* combinations, served as the reference category in a multinomial regression analysis. All explanatory variables underwent z-scaling prior to analysis (raw values below and above the mean become negative and positive, respectively). Confidence intervals of the effect estimates are represented by bars, with the effect estimates illustrated as a circle, distinct for *agr-I* and the partner *agr*-type. Non-significant terms overlap with 0 and are indicated with transparent confidence intervals.

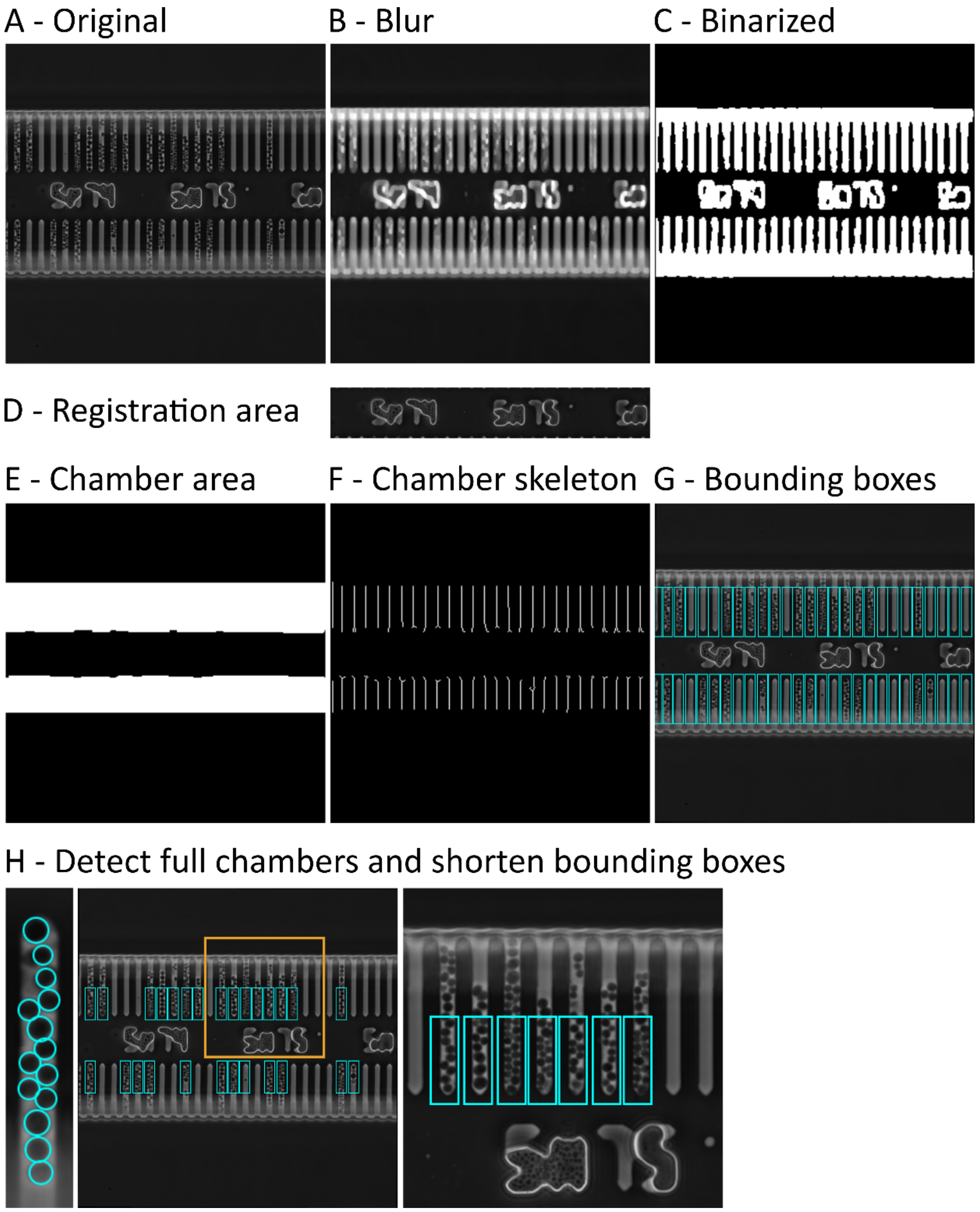


**Supplemental Figure 11**: Illustration of pre-processing of mother-machine images. The original image (A) was sequentially filtered (B) and then binarized (C). The area of the image used for registration (D) was identified by detecting the part between the chamber area (E). The binarized image (C) was cleaned up and skeletonized (F) to detect bounding boxes of each chamber (G). A Hugh transform based circle detection was performed for each chamber, chambers without any cells discarded and chamber bounding boxes shortened (H).

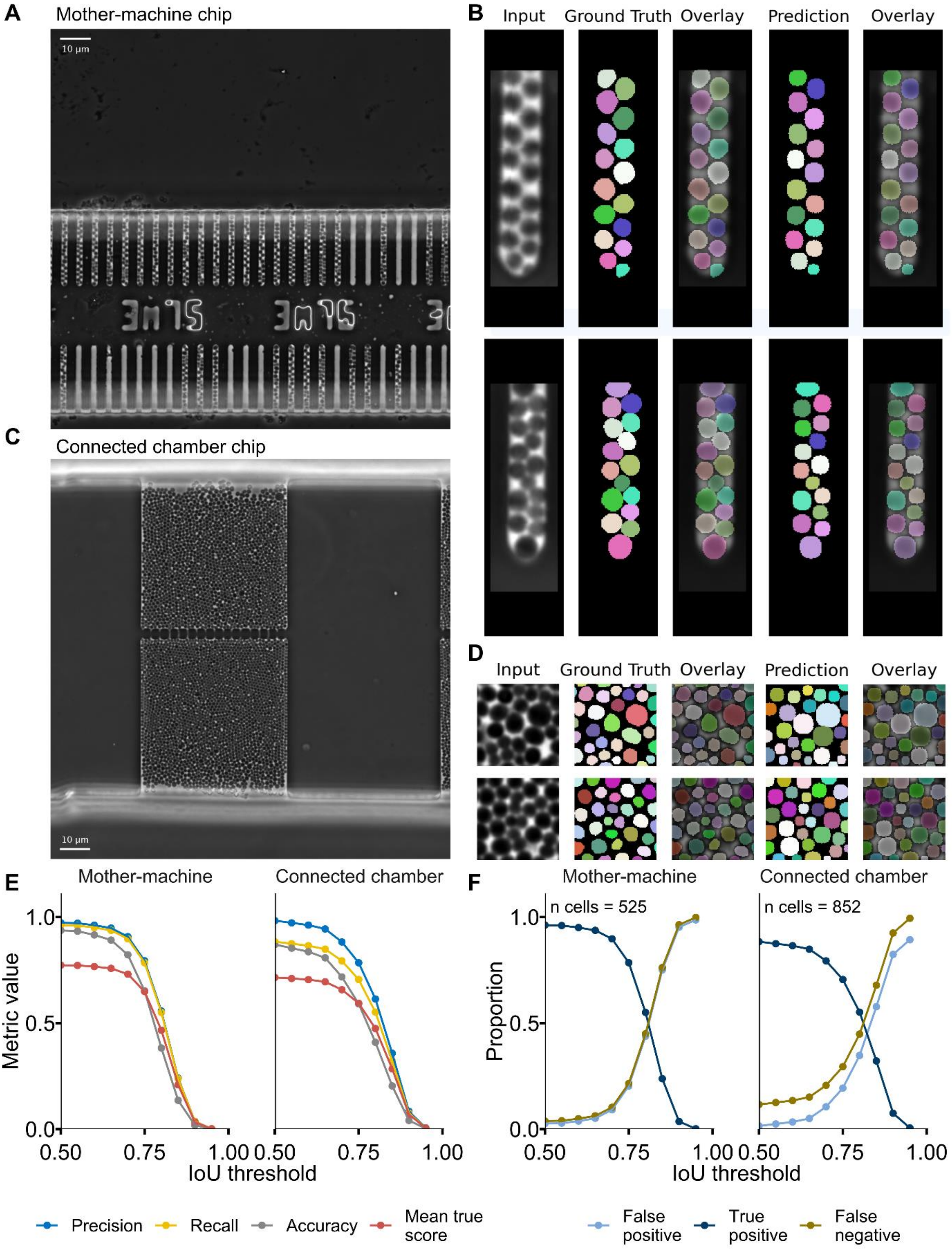


**Supplemental Figure 12: StarDist segmentation quality assessment. A.** Example image of a mother-machine microfluidic chip. Two rows of chambers can be imaged simultaneously. **B.** Example cell segmentation images of two individual mother-machine chambers using StarDist2D. **C.** Example image of a connected barrier microfluidic chip. Medium is flowing at the top and bottom and cells are embedded in the chambers. Nutrients and small molecules can diffuse through the nanochannels in the middle, but cells are retained. **D.** Example cell segmentation images of small (100 x 100 px) crops of connected chamber chips using a separately trained StarDist2D model. **E.** Segmentation model performance quantification of the mother-machine and connected chamber models. **F.** False positive, true positive and false negative rates of the segmentation models.

### Supplemental Tables

**Supplemental Table 1:** Number of observed bacterial cells for each *agr*-type and AIP concentrations used in the experiments associated with the Figures1, 2 and 3.

| | | **N cells** | |
|---|---|---|---|
| **agr type** | **AIP [nM]** | **mean per replicate** | **total** |
| agr-I | Control | 337'070 | 2'359'488 |
| agr-I | 4 nM | 321'437 | 964'312 |
| agr-I | 8 nM | 321'516 | 1'286'062 |
| agr-I | 16 nM | 305'799 | 1'223'196 |
| agr-I | 32 nM | 205'490 | 1'027'452 |
| agr-I | 64 nM | 492'798 | 2'956'790 |
| agr-I | 128 nM | 253'590 | 1'014'361 |
| agr-I | 256 nM | 240'879 | 722'637 |
| agr-II | Control | 235'713 | 1'649'994 |
| agr-II | 4 nM | 257'291 | 1'286'455 |
| agr-II | 8 nM | 191'457 | 765'828 |
| agr-II | 16 nM | 251'326 | 753'977 |
| agr-II | 32 nM | 344'393 | 1'033'179 |
| agr-II | 64 nM | 293'768 | 2'643'912 |
| agr-II | 128 nM | 320'987 | 962'960 |
| agr-II | 256 nM | 761'899 | 1'523'798 |
| agr-III | Control | 627'760 | 3'766'559 |
| agr-III | 4 nM | 525'594 | 1'576'783 |
| agr-III | 8 nM | 607'287 | 1'821'860 |
| agr-III | 16 nM | 650'440 | 1'951'319 |
| agr-III | 32 nM | 381'493 | 1'144'480 |
| agr-III | 64 nM | 438'376 | 2'630'253 |
| agr-III | 128 nM | 308'948 | 926'843 |
| agr-III | 256 nM | 305'668 | 917'003 |
| agr-IV | Control | 403'863 | 2'827'040 |
| agr-IV | 4 nM | 408'851 | 1'226'552 |
| agr-IV | 8 nM | 260'742 | 782'226 |
| agr-IV | 16 nM | 162'993 | 488'980 |
| agr-IV | 32 nM | 283'678 | 1'134'713 |
| agr-IV | 64 nM | 250'498 | 1'753'489 |
| agr-IV | 128 nM | 189'286 | 378'571 |
| agr-IV | 256 nM | 286'075 | 858'224 |
| | **Mean** | **350'843** | **1'448'728** |

**Supplemental Table 2:** Number of observed bacterial cells for mixed AIPs and concentrations used in the experiments associated with Figure 5.

| | | N cells | |
|---|---|---|---|
| **heterologous** | **AIP [nM]** | **mean per replicate** | **total** |
| mix-AIP-II | 32 nM | 158'708 | 476'123 |
| mix-AIP-II | 64 nM | 171'404 | 514'212 |
| mix-AIP-III | 32 nM | 225'849 | 677'547 |
| mix-AIP-III | 64 nM | 150'201 | 450'604 |
| mix-AIP-IV | 32 nM | 304'962 | 914'887 |
| mix-AIP-IV | 64 nM | 209'456 | 628'368 |
| | **Mean** | **203'430** | **610'290** |

**Supplemental Table 3:** Number of observed chambers for each *agr* combination tested as well as the number (N) and percentage of observation of each dynamic category for the data shown in Figure 6.

| Combination | Total observations | Category | N | Percentage |
|---|---|---|---|---|
| **agr-I vs agr-II** | 79 | stable dominance | 15 | 18.99 |
| | | initial dominance | 0 | 0.00 |
| | | switched dominance | 3 | 3.80 |
| | | delayed dual activation | 41 | 51.90 |
| | | unstable dual activation | 7 | 8.86 |
| | | stable dual activation | 13 | 16.46 |
| **agr-I vs agr-III** | 121 | stable dominance | 86 | 68.80 |
| | | initial dominance | 1 | 0.80 |
| | | switched dominance | 23 | 18.40 |
| | | delayed dual activation | 7 | 5.60 |
| | | unstable dual activation | 4 | 3.20 |
| | | stable dual activation | 4 | 3.20 |
| **agr-I vs agr-IV** | 46 | stable dominance | 11 | 23.91 |
| | | initial dominance | 2 | 4.35 |
| | | switched dominance | 3 | 6.52 |
| | | delayed dual activation | 7 | 15.22 |
| | | unstable dual activation | 2 | 4.35 |
| | | stable dual activation | 21 | 45.65 |

### Supplemental Movie legends

Applicable to all movies: Movies show phase-contrast images overlaid with fluorescent images quantifying *agr* expression via a Yellow Fluorescent Protein whose expression is controlled via the P3 promotor. The same contrast scaling has been applied to all frames of all movies per folder. Images have been registered.

##### Supplemental Movie 1

Movies relate to data shown in Fig. 1, 3 and 4 and Suppl. Fig. 2 and 5. For each *agr*-type and AIP concentration (both indicated in file name), one example movie of a single position is included. Time in HH:MM is displayed on top left, followed by indication of experimental phase (-AIP (pre), +AIP or -AIP (post)), *agr*-type and AIP concentration.

##### Supplemental Movie 2

Movies relate to data shown in Fig. 5. For each heterologous AIP type and concentration combination, one example movie of a single position is included. Time in HH:MM is displayed on top left, followed by indication of experimental phase, including type of heterologous AIP and its concentration.

##### Supplemental Movie 3

Movies relate to data shown in Fig. 6 and Suppl. Fig. 7 – 10. For each observed *agr* interaction category, one example movie of a single position is included. For the four main categories, the movies are from the same positions as shown the images in Fig. 6. Time in HH:MM is displayed on bottom left.

### Supplemental Animation legend

##### Supplemental Animation 1

Animated density histograms of the same data as shown in Fig. 1E. Histograms depict *agr* activity of bacteria observed during specific timespans (duration of 0.5 h) for each *agr*-type and AIP concentration.

### Supplemental Methods

#### Image preprocessing

To facilitate cell segmentation and tracking, we developed an image registration pipeline operating in MATLAB based on phase contrast images, which included detection of positions of filled microfluidic chambers. This pre-processing step differed slightly for mother-machine devices and connected chamber chips. For mother-machine devices, the image region containing the position labelling was detected by a multi-stage binarization algorithm and cropped out to provide a constant part for image registration. On these, multimodal estimation of the geometric transformation for translation was performed with a one-plus-one evolutionary optimization algorithm. Next, the position of chambers was detected by a sequence of filtering and binarization steps. The presence of cells was determined with a circular Hough transformation for circle detection. Only filled chambers were cropped out and copied into separate folders for further processing. The process is illustrated in Suppl. Fig. 11.

Connected chamber chip registration was performed using the same translation estimation method but on the entire downscaled and filtered image. Chamber detection was performed by detecting peaks of high intensity on the filtered image along the x and y coordinates which corresponded to the border of the area of interest.

The identified registration parameters were applied to the fluorescent images as well. Beforehand, fluorescent 16-bit images were normalized using [0.005, 0.75] • $2^{16}$ or [0.002, 0.85] • $2^{16}$, for mother-machine or connected chamber chips, respectively.

#### Plate-reader based *agr* activity

Bacteria were then inoculated at on $OD_{600nm}$ of 0.0025 in TSB containing 10 µg/ml chloramphenicol in 96 well plates, derived by diluting and washing an overnight culture. Absorbance at 600 nm and fluorescence (excitation: 500 nm, emission: 540 nm) were measured with an iD3 plate reader (Molecular Devices) in 15 min intervals over 48 h with intermittent shaking. Two-fold serial dilution (25 µl) of autoinducing peptide (AIP) in phosphate buffered saline (PBS) was performed and added to the wells once the cultures reached an OD600 of 0.1.

#### Classifying *agr* dynamics of the connected barrier chips

Mean YFP signal per community was used to binarize *agr* activity into ON and OFF state (> 100 AU). The proportion of ON state within the experimental time window of 5 to 25 h was calculated and used for *agr* dynamics classification. Only frames with at least 30 cells per chamber have been considered and only chambers with a median number of cells over time larger than 700 within the experimental time window were included. *agr* dynamics have been classified regarding the proportion of time each of the two interacting communities are in activated *agr* state and the relative activity of these two.

Dominance was defined as minimally 40% unequal ON state between the two interacting *agr*-types, maximally 10% simultaneous ON state and either of the communities being ON for at least 60% and the other being ON maximally for 15%.

Initial dominance was defined as maximally 15% of simultaneous ON state, minimally 30% unequal ON state and one of the communities being ON for less than 60% and being ON already in the first 40% of observed time.

Switched dominance was defined as at least 30% unequal ON state among the interacting *agr*-types and either of the communities being ON for minimally 15% and maximally 80% and either of them being ON in the early phase of the experiment (first 40% of experimental window).

Delayed dual activation was defined as intermediate proportion of unequal ON state among the interacting *agr*-types (> 30%) and at least 10% of simultaneous ON state. Either one of the communities was required to be ON for at least 40% of the time and of the other more than 65%.

Unstable dual activation was defined as at least 30% of simultaneous OFF state and either community being in an ON state for more than 20%.

Stable dual activation was defined as at least 40% simultaneous ON state and maximally 40% of unequal ON state.
