## Supplementary figures and images for "Single-cell approach dissecting *agr* quorum sensing dynamics in *Staphylococcus aureus*"

### Supplemental Animation 1

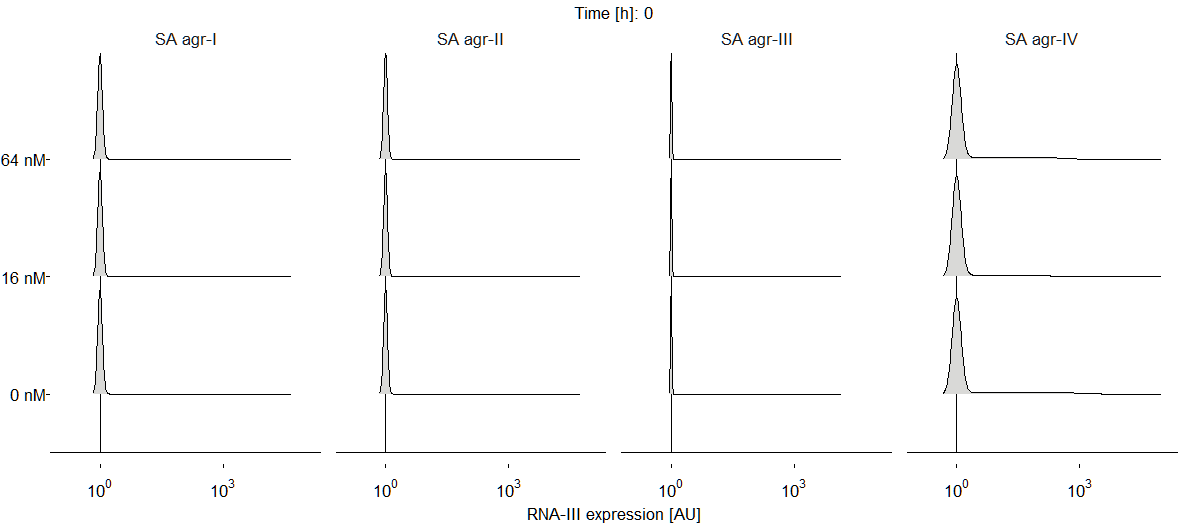
